# Phytoplankton community composition in the oligotrophic Argo Basin of the eastern Indian Ocean

**DOI:** 10.1101/2025.08.14.670423

**Authors:** Karen E. Selph, Robert H. Lampe, Natalia Yingling, Ria Bhabu, Sven A. Kranz, Andrew E. Allen, Michael R. Landry

**Author notes:** Corresponding author E-mail address (K.E. Selph).

## Abstract

Phytoplankton community composition during austral summer 2022 in the Argo Abyssal Plain (Argo Basin), a 5000-m deep area northwest of the Australian continent in the eastern Indian Ocean, is described in detail, including phytoplankton abundance, biomass, size structure, taxonomic identifications through DNA and pigment analyses, as well as the percent of functional mixotrophs. The region was characterized by warm (up to 30.5°C), stratified, oligotrophic (nitrogen-limited) waters, with integrated euphotic zone (EZ) chlorophyll a (CHL*a*) of 13 mg m^-2^. The EZ mean CHL*a* was low in the upper layer (0.085 µg L^-1^) and 0.32 µg L^-1^ at the pronounced deep CHL*a* maxima. EZ-integrated phytoplankton carbon averaged 1229 mg C m^-2^. *Prochlorococcus* was the dominant taxon throughout the EZ, but the lower EZ had ∼4-times more eukaryotic carbon biomass than the upper EZ, along with a distinct community. In the upper EZ, prymnesiophytes, dinoflagellates and prasinophyte taxa without prasinoxanthin had the highest contributions to monovinyl chlorophyll a (MV-CHL*a*). In the lower EZ the community was more diverse, with prymnesiophytes, dinoflagellates, prasinophyte taxa with prasinoxanthin, pelagophytes, and cryptophytes all comprising significant contributions to MV-CHL*a*. Diatoms were a minor part of the community. In the upper EZ, a higher percent of the community showed mixotrophy (35-84%) relative to the lower EZ (30-51%). Although a low abundance, nitrogen-fixing organisms (symbionts of diatoms and cyanobacteria taxa) were ubiquitous. Overall, the community was similar to that found at the Hawaii Ocean Time-series site and the central Gulf of Mexico.

## 1. Introduction

Phytoplankton community structure is shaped by light, nutrients, temperature and ecological interactions, such as mortality (grazers, viruses) and competition, leading to the expectation that similar oceanic environments will harbor similar taxa. In surface waters of low-nutrient regions of the open oceans from temperate to tropical zones, prokaryotes, especially *Prochlorococcus*, and picoeukaryotes (<2 µm diameter cells) represent about half of the phytoplankton biomass (Raven, 1998; Partensky et al., 1999; Moon-van der Staay et al., 2001; Buitenhuis et al., 2012). The picoeukaryote taxa are more diverse, with prymnesiophytes and prasinophytes making up a large proportion (Kirkham et al., 2013; Cabello et al., 2016). The rest remaining cells with photosynthetic functionality, nano- and microeukaryotes (2-20 µm and 20-200 µm cells, respectively), are mostly dinoflagellates, with some larger prymnesiophytes (Taylor and Landry, 2018). Deeper waters, closer to the nitracline, usually show a similar community structure, with the addition of pelagophytes and cryptophytes, which are better adapted to the lower light quality and quantity at depth and to higher nutrient conditions (Stomp et al., 2007; Guérin et al., 2022; Richardson, 2022).

Despite the similarities of the phytoplankton taxa residing in low nutrient waters, they represent groups occupying diverse ecological niches. For example, some are considered strict autotrophs, while others are mixotrophs (Stoecker, 1999; Anderson et al., 2018; Edwards et al., 2023). Some thrive under high light/low nutrient conditions, whereas others require higher nutrient levels but can live in low light environments (Latasa et al., 2023; Eckmann et al., 2024). Additionally, some lack the ability to use nitrate and instead rely on ammonium (Moore et al., 2002), while others are capable of nitrogen fixation or form symbiotic relationships with nitrogen-fixing cyanobacteria (Mague et al., 1977; Foster et al., 2011). Since phytoplankton community composition drives higher trophic level community composition (Irwin et al., 2006; Sommer et al., 2018; Décima, 2022), resolving the distributions of these foundational organisms in habitats of interest contributes to modeling efforts for fisheries and climate change.

Waters overlying the Argo Abyssal Plain (hereafter, Argo Basin) northwest of the Australian continent and south of Java, Indonesia in the eastern Indian Ocean, comprise a poorly studied oligotrophic system. Not et al. (2008) and Schlüter et al. (2011) conducted transects that ended near this area, however their stations were close enough to land to be influenced by coastal processes, whereas the land-remote Argo Basin is mainly affected by mesoscale features, such as transient eddies (Kehinde et al., 2023). It is also an important fishery area, as the only place where Southern Bluefin Tuna (SBT) are known to spawn, with their larvae relying on the local prey field for survival (Matsuura et al., 1997; Landry et al., this issue). In a warming world, determining current characteristics and vulnerabilities to change is critical for understanding future stock recruitment as well as other effects on the global carbon cycle (Cetinić et al., 2024).

As part of the BLOOFINZ (Bluefin Larvae in Oligotrophic Ocean Foodwebs, Investigations of Nitrogen to Zooplankton) project, we investigated the Argo Basin and surrounding areas in the austral summer during the peak time of SBT spawning (Landry et al., this issue). Here, we assess phytoplankton abundance, biomass and size structure from depth profiles of phytoplankton distributions covering the euphotic zone (EZ), as well as surface transects through the area. Results are compared to a previous study by the same methods in the Gulf of Mexico (GoM), a spawning habitat for the Atlantic Bluefin Tuna (Selph et al., 2021; Gerard et al., 2022). For the Argo Basin, we add DNA sequence analyses of dominant taxa, as well as assessments of what percent of functional mixotrophs exist among the chlorophyll-bearing protists.

## 2. Materials and methods

### 2.1. Study area and general sampling scheme

The BLOOFINZ study was conducted in the Argo Basin off NW Australia aboard the R/V *Roger Revelle* on cruise RR2201 from 26 January to 6 March 2022 (Landry et al., this issue). We sampled phytoplankton populations in open-ocean waters using a 12-place rosette of Niskin bottles, Niskin-X bottles hung on hydrowire, and through the ship’s underway flowing seawater system (∼5-m inlet depth). Each set of stations was part of 3-5 d Lagrangian experiment (hereafter, “cycle”) of daily-repeated sampling and incubation activities following a satellite-tracked drift array with 3×1-m holey sock drogue centered at 15 m (Landry et al., 2009; Landry et al., this issue). The locations of the initial CTD cast (02:00) for each cycle experiment was determined from physical and biological surveys and satellite imagery and marked with the deployments of free-drifting in situ sediment trap and incubation bottle arrays. Concurrently and between cycles, underway sampling was done using the ship’s underway seawater system to collect surface seawater samples.

### 2.2. Environmental measurements

The rosette-mounted CTD (Seabird Scientific 911) had 12 Niskin bottles and sensors for temperature, conductivity, chlorophyll (CHL*a*) fluorescence and oxygen. Each cast sampled 6-8 depths in the EZ (to the 1% incident irradiance depth). The CHL*a* profiling sensor (Wet Labs FLRTD-1156) was calibrated with discrete samples collected from the Niskin bottles, which were filtered onto GF/Fs, extracted in 90% acetone for 24 h (dark, −20°C), then analyzed on a Turner Designs Model 10 fluorometer using the acidification method (Strickland and Parsons, 1972). A rosette-frame mounted photosynthetically active radiation (PAR) sensor, Biospherical Model QSP200L4S, with a 2-pi light sensor was deployed daily after sunrise to determine in situ irradiance profiles at each station.

Nutrient samples (nitrate+nitrite, ammonium, phosphate, and silicic acid) were taken by 0.2-µm filtration directly from the Niskin bottles, frozen, and analyzed on shore by the Ocean Data Facility (Scripps Institution of Oceanography) using an auto-analyzer (Becker et al., 2019). Fluorometric CHL*a* was also analyzed on shipboard from 250-mL samples processed as described above for calibrating the profiling CHL*a* sensor. Phytoplankton community composition was determined by flow cytometry (FCM), high-performance liquid chromatography (HPLC), epifluorescence microscopy (EPI-MICRO), inverted microscopy (INVERT-MICRO) and an Imaging FlowCytoBot (IFCB). Samples from the surface mixed layer and the deep CHL*a* maximum (DCM) were analyzed for DNA-based abundances of the phytoplankton community. These methods are described in the following subsections and abbreviations used are defined in Table 1. When data are reported as means, error is ± 1 standard error of the mean.

**Table 1.** Abbreviations used in the text.

|  |  |  |
| --- | --- | --- |
| 1184 | **Table 1.** Abbreviations used in the text. |  |
| 1185 |  |  |
| 1186 | Allophycocyanin (cryptophyte pigment) | ALLO |
| 1187 | Amplicon sequence variant | ASV |
| 1188 | Autotrophic eukaryotes | A-EUK |
| 1189 | Autotrophic dinoflagellates | A-DINO |
| 1190 | 19'-butanoyloxyfucoxanthin (pelagophyte pigment) | BUT |
| 1191 | Cryptophytes | CRYPT |
| 1192 | Chlorophyll a | CHL<sub>a</sub> |
| 1193 | Diatoms | DIAT |
| 1194 | Divinyl chlorophyll a (*Prochlorococcus* pigment) | DV-CHL<sub>a</sub> |
| 1195 | Euphotic zone (surface to 1% incident irradiance) | EZ |
| 1196 | Flow cytometry | FCM |
| 1197 | Fucoxanthin (diatom pigment) | FUCO |
| 1198 | 19'-hexanoyloxyfucoxanthin (prymnesiophyte pigment) | HEX |
| 1199 | High-performance liquid chromatography | HPLC |
| 1200 | Imaging FlowCytoBot | IFCB |
| 1201 | Inverted microscopy | INVERT-MICRO |
| 1202 | Lutein (prasinophyte pigment) | LUT |
| 1203 | Epifluorescence microscopy | EPI-MICRO |
| 1204 | Eukaryotes | EUK |
| 1205 | Microeukaryotes (>20 µm cell diameter) | MICRO-EUKS |
| 1206 | Monovinyl chlorophyll a (SYN and all other PEUK) | MV-CHL<sub>a</sub> |
| 1207 | Monovinyl chlorophyll b (prasinophyte pigment) | MV-CHL<sub>b</sub> |
| 1208 | Mixotrophic Eukaryotes (have acidic vacuoles) | MEUK |
| 1209 | Nano-eukaryotes (2-20 µm cell diameter) | NANO-EUKS |
| 1210 | Neoxanthin (prasinophyte pigment) | NEOX |
| 1211 | Pelagophytes | PELAG |
| 1212 | Peridinin (autotrophic dinoflagellate pigment) | PER |
| 1213 | Photosynthetic eukaryotes | PEUK |
| 1214 | Picoeukaryotes (0.8-2µm cell diameter) | PICO-EUKS |
| 1215 | Prasinoxanthin (prasinophyte group 3 pigment) | PRAS |
| 1216 | Prasinophytes group 1 | PRAS-1 |
| 1217 | Prasinophytes group 3 | PRAS-3 |
| 1218 | *Prochlorococcus* | PRO |
| 1219 | Prymnesiophytes (= haptophytes) | PRYM |
| 1220 | *Synechococcus* | SYN |
| 1221 | Total Chlorophyll a (MV- plus DV-CHL<sub>a</sub> from HPLC) | TCHL<sub>a</sub> |
| 1222 | Violaxanthin (prasinophyte pigment) | VIOL |
| 1223 | Zeaxanthin (found in many groups, photoprotective) | ZEAX |

### 2.3. Phytoplankton from flow cytometry

Samples (1 mL) for determination of microbial populations were analyzed on shipboard using a Beckman Coulter CytoFlex S flow cytometer. Both live (unpreserved) and dead (preserved) samples were collected from the same Niskin bottles. No significant difference was found for prokaryotes between live and preserved samples (i.e., ≤10% between *Prochlorococcus* determinations); therefore most of the data presented were from preserved samples. Preserved (0.5% paraformaldehyde, v/v, final) samples were stained with Hoechst 34580 (DNA stain), incubated for 1 h (room temperature, dark), then analyzed at 50 µL min^-1^ for 10 min (500 µL sample volume), exciting with 375 nm (em 450±45 nm, blue fluorescence for DNA), 488 nm (em 690±50 nm, red fluorescence for CHLa), 561 nm (em 585±42 nm, orange fluorescence for phycoerythrin), and collecting 405 nm and 488 nm light scatter (forward and 90° side scatter) signals. Details of gating schemes and sample analyses are as in Selph (2021). A subset of live samples was stained with LysoTracker Green (green acidic vacuole stain; Rose et al., 2004) to determine abundances of mixotrophic eukaryotic plankton (MEUK, CHL*a*-bearing cells with digestive vacuoles indicating current phagotrophy). Details of LysoTracker Green staining and subsequent sample processing are in Selph et al. (this issue); however, the signal from these cells was generated by the 488 nm laser (em 525±40 nm, green fluorescence). Populations identified from these analyses were *Prochlorococcus* (PRO), *Synechococcus* (SYN), photosynthetic and mixotrophic eukaryotes (PEUK and MEUK, respectively).

Cell abundances from FCM were converted to estimated carbon biomass using 32 and 101 fg C cell^-1^ for PRO and SYN, respectively (Garrison et al., 2000). Picoeukaryotes, <2-µm cell diameter MEUK and PEUK, were estimated to be 320 fg C cell^-1^ using Menden-Deuer and Lessard’s (2000) equation (pg C cell^-1^ = 0.216 x BV^0.939^) and estimating their mean biovolume (BV) as that of 0.8-1.9 µm diameter cells. Larger cell biomass (>2-µm) was derived from microscopy (Section 2.4, detailed below).

### 2.4. Pigment analyses and taxonomic assignments

Samples for HPLC pigment analyses (2.3 L) were collected from Niskin bottles, filtered onto GF/F filters, then flash frozen in liquid nitrogen and stored at −80°C. Samples were extracted (2 h in 3 mL 100% methanol), sonicated, clarified by GF/F filtration, and analyzed on an HPLC Agilent Technologies 1200 instrument at the Institut de la Mer de Villefranche, Laboratorie d’Oceanographie de Villefranche (Ras et al., 2008).

Pigments used in taxonomic classifications were divinyl chlorophyll *a* (DV-CHL*a*), monovinyl chlorophyll a (MV-CHL*a*), monovinyl chlorophyll *b* (MV-CHL*b*), peridinin (PER), 19’-hexanoyloxyfucoxanthin (HEX), 19’-butanoyloxyfucoxanthin (BUT), prasinoxanthin (PRAS), fucoxanthin (FUCO), neoxanthin (NEO), lutein (LUT), violaxanthin (VIOL) and alloxanthin (ALLO). DV-CHL*a* is only found in PRO, whereas MV-CHL*a* is found in all other taxa. Given the lack of a unique pigment to discriminate SYN from other taxa, and the good correlation (r^2^ = 0.89) between the sum of red fluorescence (flow cytometry indicator of CHL*a*) and *T*CHL*a* (sum of DV-CHL*a* and MV-CHL*a* from HPLC, Supp. Fig. S1), we used the relative ratios of FCM-derived abundances of SYN and PRO compared to DV-CHL*a* to determine the amount of MV-CHL*a* to assign to SYN (Selph et al., 2021). The remaining MV-CHL*a* was partitioned into taxonomic groups using the chemotaxonomic R-program phytoclass (v. 2.0.0; Hayward et al., 2023), with final pigment ratios shown in Supp. Table S1. For the analyses, samples were divided into shallow (<50 m) and deep (≥50 m) bins, as it was evident that those depth horizons comprised somewhat different community compositions. The taxonomic groups identified were prymnesiophytes (PRYM), pelagophytes (PELAG), diatoms (DIAT), autotrophic dinoflagellates (A-DINO), cryptophytes (CRYPT), and 2 groups of chlorophytes. The chlorophyte groups are comprised of prasinophyte group 3 with PRAS (PRAS-3), which includes *Ostreococcus* and *Bathycoccu*s, and prasinophyte group 1 (PRAS-1), which has the basic chlorophyte pigments but no PRAS. Abbreviations for pigments and taxonomic groups are defined in Table 1.

### 2.5. Microscopy for autotrophic biomass

Samples (500 mL) for autotrophic nano- and micro-plankton abundance and biomass were prepared for epifluorescence microscopy (EPI-MICRO) after gravity collection from Niskin bottles using silicone tubing, preservation with sequential additions of alkaline Lugol’s, formaldehyde, and sodium thiosulfate, staining with proflavine (0.33% w/v), and gentle vacuum filtration (<100 mm Hg) onto 0.8 µm (50 mL) and 8 µm (450 mL) black PCTE 25 mm filters supported by 10 µm nylon filters (Sherr and Sherr, 1993; Taylor et al., 2015). When ∼10 mL of sample was left in the filter tower, the samples were stained with 4’,6-diamidino-2-phenylindole (DAPI) for 2 min and filtered to dryness. Black filters were mounted on glass slides, with a small drop of immersion oil on the filter then capped with a cover slip. All slides were frozen (−80°C) until shore-based image analyses. Slide imaging was done using an Olympus Microscope DP72 Camera mounted on an Olympus BX51 fluorescence microscope. Images were captured under 20x magnification (8-µm filters) and 60x magnification (0.8 µm filters), and analyzed using ImageJ software (Yingling et al., this issue).

Diatoms and dinoflagellates were also enumerated using inverted microscopy (INVERT-MICRO) from acid Lugol’s (5% final solution) preserved samples (250 mL), with a glass cover slip added to the solution to help ensure silica saturation to prevent diatom frustule dissolution (Throndsen, 1978). Two samples per cycle were analyzed after pooling samples from the upper EZ (0-40 m) and the lower EZ (40 m to base of the EZ). Of these, 320-480 mL of sample was gravity settled for 24 h and enumerated and imaged with a Zeiss AxioVert 200M inverted microscope and Zeiss AxioCam MRm CCD camera (Utermöhl technique; Lund et al., 1958).

Biovolume estimates were derived from feret length, cell area and width for EPI-MICRO images (Yingling et al., 2025), and cell length and width measurements for INVERT-MICRO images. These dimensions were converted to biovolumes, then carbon equivalents using the Menden-Deuer and Lessard (2000) equations: pg C cell^-1^ = 0.216 x BV^0.939^ for non-diatoms and pg C cell^-1^ = 0.288 x BV^0.811^ for diatoms.

### 2.6. Underway throughflow sampling for Imaging FlowCytobot (IFCB)

Imaging flow cytometry of the eukaryotic phytoplankton community (∼8 to 150 µm) was performed with an imaging flowcytobot (IFCB) (Olson and Sosik, 2007). The IFCB was connected to the ship’s underway seawater system that used a Grace Husky 1050e diaphragm pump in accordance with best practices (Cetinić et al., 2016). During most of the cycles, the IFCB automatically collected 5-mL samples approximately every 25 min. Imaging was triggered by CHL*a* fluorescence.

Biovolume estimates for IFCB images were determined by the automated distance map algorithm described by Moberg and Sosik (2012). Carbon estimates were calculated from the computed biovolumes using the Menden-Deuer and Lessard (2000) equations for diatoms and non-diatoms.

### 2.7. DNA-based characterization of the eukaryotic phytoplankton community

For DNA, seawater was filtered onto 0.22 µm Sterivex-GP filters (Cat. no. SVGP01050, Millipore) that were immediately sealed with luer-lock plug and Hemato-Seal™ tube sealant, wrapped in aluminum foil, and flash-frozen in liquid nitrogen. For samples from the CTD rosette, between 1.5 L and 4 L (mean = 3.1 L) was filtered from either 10 m or the CHL*a* maximum. At Argo stations, 2 L samples from each depth were obtained from Niskin-X bottles on a Kevlar line configured for trace metal clean sampling. Underway samples (1-4 L, mean = 3.6 L) were also collected both during cycles and transects from the ship’s underway seawater system and towed surface fish. Filters were stored at −80°C until extraction.

DNA was extracted with the Macherey-Nagel NucleoMag Plant Kit for DNA purification (Cat. No 744400) on an Eppendorf epMotion 5075TMX. At the start of DNA extraction during the addition of lysis buffer, 5.78 or 6.70 ng of *Schizosaccharomyces pombe* genomic DNA and 7.78 or 8.95 ng of *Thermus thermophilus* genomic DNA were added to each sample as an internal standard (Lin et al., 2019). DNA was then assessed on a 1.8% agarose gel.

Amplicon libraries targeting the V4-V5 region of the 16S rRNA gene or the V4 region of the 18S rRNA gene were generated via a one-step PCR using the Azura TruFi DNA Polymerase PCR kit. For 16S, the 525F-Y (5’-GTG YCA GCM GCC GCG GTA A-3’) and 926R (5’-CCG YCA ATT YMT TTR AGT TT-3’) primer sets were used (Parada et al., 2016). For 18S, the V4F (CCAGCASCYGCGGTAATTCC) and V4RB (ACTTTCGTTCTTGATYR) primers modified from Berdjeb et al. (2018) were used. Each reaction was performed with an initial denaturing step at 95°C for 1 min, followed by 30 cycles of 95°C for 15 s, 56°C for 15 s, and 72°C for 30 s. To confirm amplification, 2.5 µL of each PCR reaction was run on a 1.8% agarose gel. PCR products were purified using a 1x volume of Cytiva Sera-Mag Select beads. PCR quantification was performed using Invitrogen Quant-iT PicoGreen dsDNA Assay kit. Samples were then pooled in equal proportions followed by a 0.8x magnetic bead PCR purification. The final pools were then evaluated on an Agilent 2200 TapeStation and quantified with Qubit HS dsDNA kit. Sequencing was performed at the gCore at the University of California San Diego on an Illumina NextSeq 2000 (2 × 300 bp) with a 25% PhiX spike-in. Mock community samples were prepared and added to the pool alongside the environmental samples here (Lampe et al., 2025).

Amplicons were analyzed with QIIME2 v2024.5 (Bolyen et al., 2019). Briefly, demultiplexed paired-end reads were trimmed to remove adapter and primer sequences with cutadapt (Martin, 2011). Trimmed reads were then denoised with DADA2 to produce amplicon sequence variants (ASVs; Callahan et al., 2016). Taxonomic annotation of ASVs was conducted with the q2-feature-classifier classify-sklearn naïve-bayes classifier (Bokulich et al., 2018) and either the SILVA (Release 138) or PR^2^ database (v. 5.0.0) (Pruesse et al., 2007; Guillou et al., 2013). Phytoplankton taxa were then filtered for subsequent analysis based on lineages that are known photoautotrophs, resulting in the following groupings that align with other measurements – chlorophyll-bearing dinoflagellates (A-DINO), CRYPT, DIAT, PELAG, PRYM, PRAS, PRO, SYN and OTHER. The DNA-based PRAS group is comprised of both PRAS-1 and PRAS-3.

Alpha and beta diversity metrics were computed with the vegan package in R (Oksanen et al., 2007). To estimate absolute abundances of ASVs, recovery of the aforementioned internal standards was used (Lin et al., 2019). For each ASV within each sample, the number of reads was divided by the ratio of *T. thermophilus* or *S. pombe* reads to the number of rRNA copies added. The total number of copies was then normalized to the volume filtered for each sample to estimate rRNA copies L^-1^. Raw sequence data have been deposited in the NCBI Sequence Read Archive under BioProject accession number PRJNA1281638.

## 3. Results

### 3.1. Study area characteristics

The study area was entirely in the open-ocean waters of the Argo Basin (Fig. 1). Stations were occupied from 3 Feb – 2 March 2022. Cycle 1 stations were occupied after a tropical storm went through the region and had the deepest mixed layers (mean 36.2 ± 4.6 m), whereas subsequent cycles were more stratified and had mixed layers averaging 11 ± 1.4 m (Table 2). Mixed-layer water temperatures ranged from 28.45 to 30.55°C, showing a general trend of warming from Cycles 1 to 4, but cooling slightly to 29.40°C for transect stations in the central Argo Basin. Except for the first station (CTD 11) of Cycle 1, where nutrient samples were slightly enhanced due to storm mixing (0.09 µM nitrate, 0.04 µM ammonium, 0.16 µM phosphate), nutrient concentrations in surface waters were very low (means of 0.01 ± 0.002 µM nitrate, 0.06 ± 0.05 µM ammonium, 0.07 ± 0.01 µM phosphate). CHL*a* concentrations in the mixed layer averaged 0.08 ± 0.004 µg L^-1^, and all stations were characterized by deep DCMs, with the maximum ranging from 60 to 94 m and profiling fluorometer CHL*a* averaging 0.58 ± 0.02 µg L^-1^ (Table 2; Kranz et al., this issue). These physical characteristics also affected productivity patterns observed during the cruise, as described in Kranz et al. (this issue), where net community production, net and gross primary productivity, and photophysiology were closely tied to episodic mixing events and restratification (Kranz et al., this issue).

**Figure 1.**
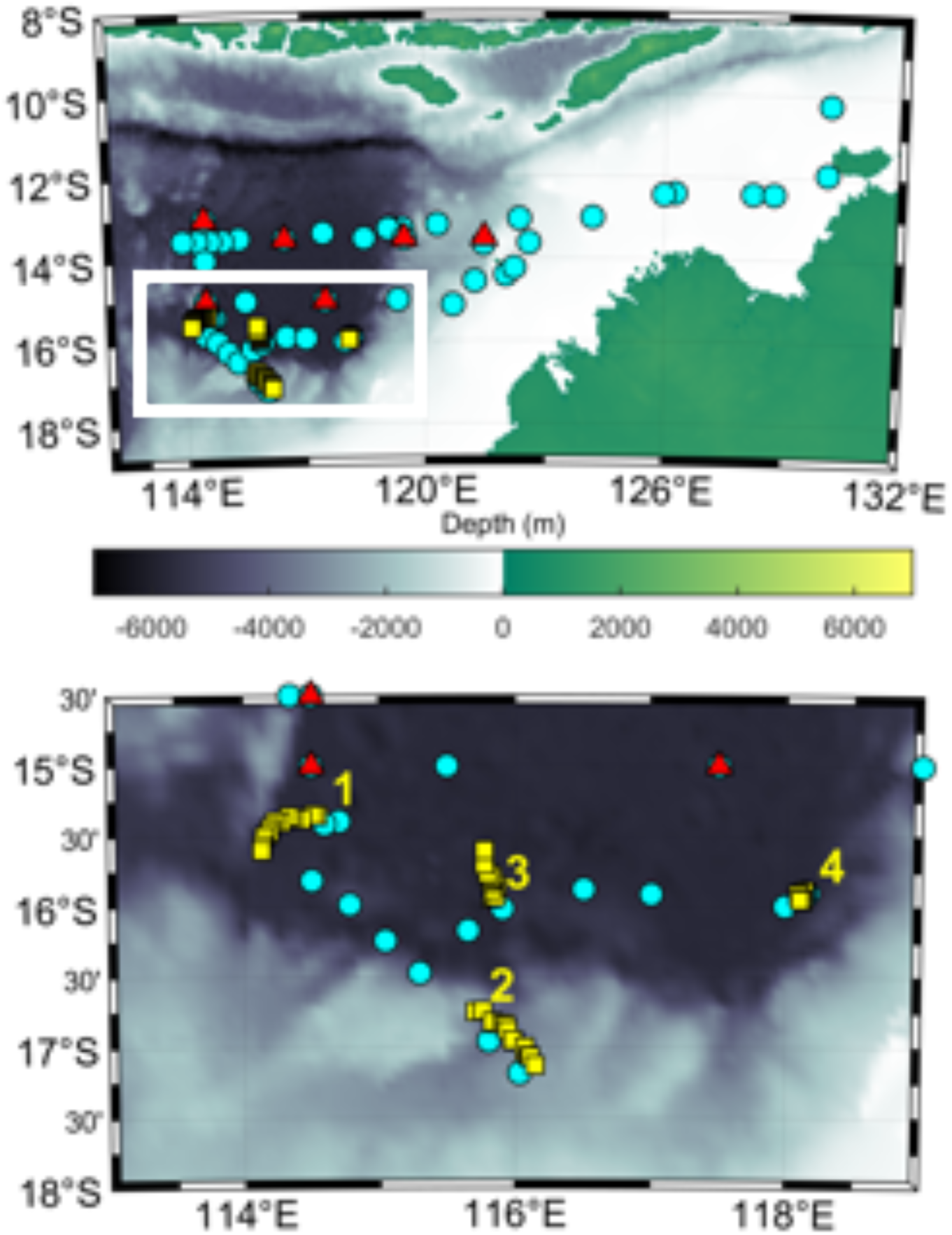
Map of study area (courtesy of M. Stukel). A) General study area showing NW Australia and parts of Indonesia, with white box showing area for lower figure; B) study area (white box in A), showing Cycle locations (1-4), a subset of Argo stations, and a subset of underway (surface sample) stations. Symbols are: YELLOW – Cycle 1-4 stations, RED – Argo TM stations, BLUE – underway (inline ship or towed fish) stations.

**Table 2.**
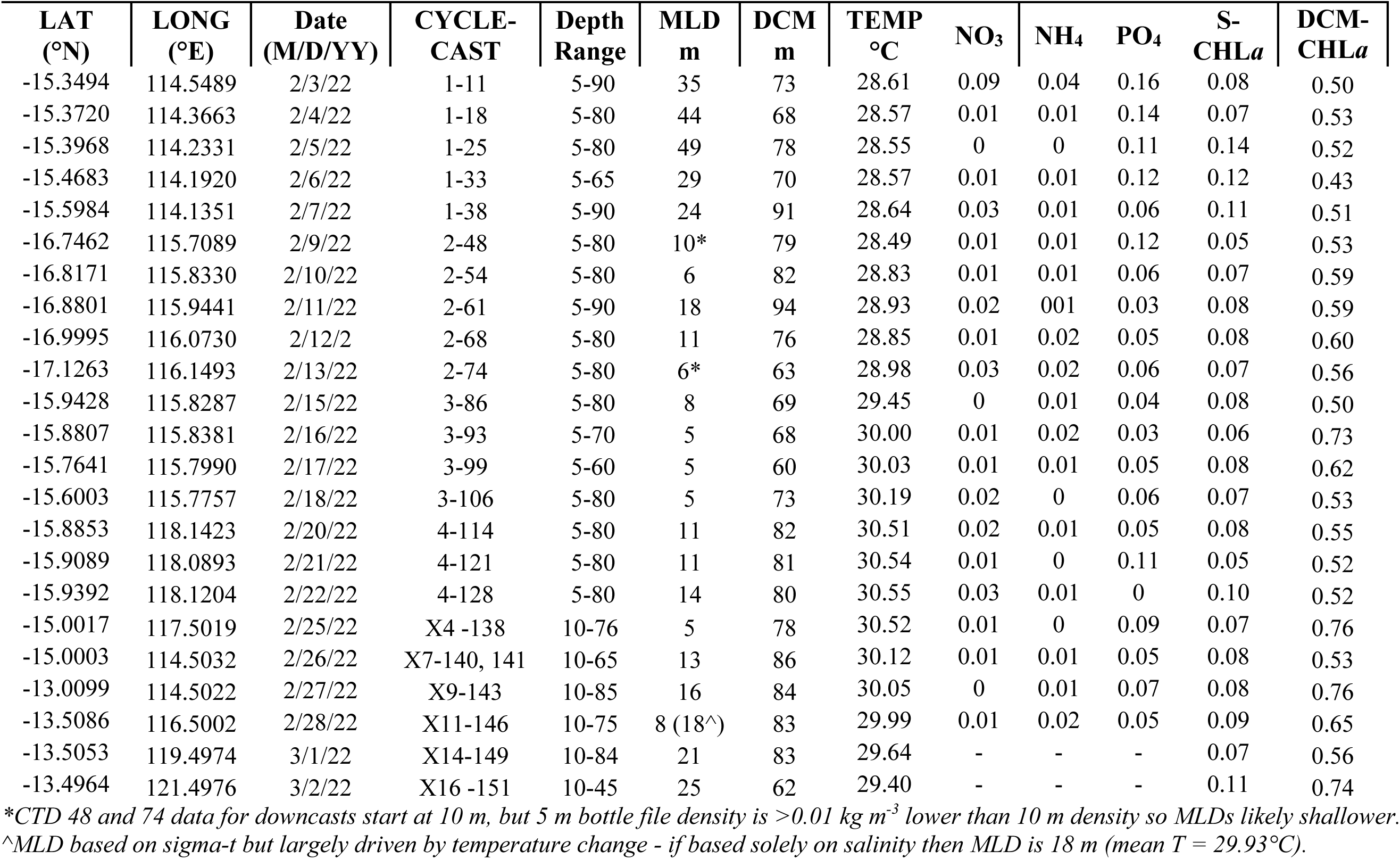
Station locations, with subsurface properties for the Niskin-X samples coming from the indicated CTD cast (G#-CTD#). Depths (m) are ranges of discrete samples, mixed layer depth (MLD), and deep chlorophyll maximum (DCM). Temperature (°C), nutrients (µM) are nitrate (NO_3_), ammonium (NH_4_), and phosphate (PO_4_); Chlorophyll *a* (CHL*a*, µg L^-1^) are mixed layer (S) and DCM values.

| LAT<br>(°N) | LONG<br>(°E) | Date<br>(M/D/YY) | CYCLE-<br>CAST | Depth<br>Range | MLD<br>m | DCM<br>m | TEMP<br>°C | NO<sub>3</sub> | NH<sub>4</sub> | PO<sub>4</sub> | S-CHL*a* | DCM-CHL*a* |
| --- | --- | --- | --- | --- | --- | --- | --- | --- | --- | --- | --- | --- |
| -15.3494 | 114.5489 | 2/3/22 | 1-11 | 5-90 | 35 | 73 | 28.61 | 0.09 | 0.04 | 0.16 | 0.08 | 0.50 |
| -15.3720 | 114.3663 | 2/4/22 | 1-18 | 5-80 | 44 | 68 | 28.57 | 0.01 | 0.01 | 0.14 | 0.07 | 0.53 |
| -15.3968 | 114.2331 | 2/5/22 | 1-25 | 5-80 | 49 | 78 | 28.55 | 0 | 0 | 0.11 | 0.14 | 0.52 |
| -15.4683 | 114.1920 | 2/6/22 | 1-33 | 5-65 | 29 | 70 | 28.57 | 0.01 | 0.01 | 0.12 | 0.12 | 0.43 |
| -15.5984 | 114.1351 | 2/7/22 | 1-38 | 5-90 | 24 | 91 | 28.64 | 0.03 | 0.01 | 0.06 | 0.11 | 0.51 |
| -16.7462 | 115.7089 | 2/9/22 | 2-48 | 5-80 | 10* | 79 | 28.49 | 0.01 | 0.01 | 0.12 | 0.05 | 0.53 |
| -16.8171 | 115.8330 | 2/10/22 | 2-54 | 5-80 | 6 | 82 | 28.83 | 0.01 | 0.01 | 0.06 | 0.07 | 0.59 |
| -16.8801 | 115.9441 | 2/11/22 | 2-61 | 5-90 | 18 | 94 | 28.93 | 0.02 | 0.01 | 0.03 | 0.08 | 0.59 |
| -16.9995 | 116.0730 | 2/12/22 | 2-68 | 5-80 | 11 | 76 | 28.85 | 0.01 | 0.02 | 0.05 | 0.08 | 0.60 |
| -17.1263 | 116.1493 | 2/13/22 | 2-74 | 5-80 | 6* | 63 | 28.98 | 0.03 | 0.02 | 0.06 | 0.07 | 0.56 |
| -15.9428 | 115.8287 | 2/15/22 | 3-86 | 5-80 | 8 | 69 | 29.45 | 0 | 0.01 | 0.04 | 0.08 | 0.50 |
| -15.8807 | 115.8381 | 2/16/22 | 3-93 | 5-70 | 5 | 68 | 30.00 | 0.01 | 0.02 | 0.03 | 0.06 | 0.73 |
| -15.7641 | 115.7990 | 2/17/22 | 3-99 | 5-60 | 5 | 60 | 30.03 | 0.01 | 0.01 | 0.05 | 0.08 | 0.62 |
| -15.6003 | 115.7757 | 2/18/22 | 3-106 | 5-80 | 5 | 73 | 30.19 | 0.02 | 0 | 0.06 | 0.07 | 0.53 |
| -15.8853 | 118.1423 | 2/20/22 | 4-114 | 5-80 | 11 | 82 | 30.51 | 0.02 | 0.01 | 0.05 | 0.08 | 0.55 |
| -15.9089 | 118.0893 | 2/21/22 | 4-121 | 5-80 | 11 | 81 | 30.54 | 0.01 | 0 | 0.11 | 0.05 | 0.52 |
| -15.9392 | 118.1204 | 2/22/22 | 4-128 | 5-80 | 14 | 80 | 30.55 | 0.03 | 0.01 | 0 | 0.10 | 0.52 |
| -15.0017 | 117.5019 | 2/25/22 | X4 -138 | 10-76 | 5 | 78 | 30.52 | 0.01 | 0 | 0.09 | 0.07 | 0.76 |
| -15.0003 | 114.5032 | 2/26/22 | X7-140, 141 | 10-65 | 13 | 86 | 30.12 | 0.01 | 0.01 | 0.05 | 0.08 | 0.53 |
| -13.0099 | 114.5022 | 2/27/22 | X9-143 | 10-85 | 16 | 84 | 30.05 | 0 | 0.01 | 0.07 | 0.08 | 0.76 |
| -13.5086 | 116.5002 | 2/28/22 | X11-146 | 10-75 | 8 (18^) | 83 | 29.99 | 0.01 | 0.02 | 0.05 | 0.09 | 0.65 |
| -13.5053 | 119.4974 | 3/1/22 | X14-149 | 10-84 | 21 | 83 | 29.64 | - | - | - | 0.07 | 0.56 |
| -13.4964 | 121.4976 | 3/2/22 | X16 -151 | 10-45 | 25 | 62 | 29.40 | - | - | - | 0.11 | 0.74 |

\*CTD 48 and 74 data for downcasts start at 10 m, but 5 m bottle file density is >0.01 kg m<sup>-3</sup> lower than 10 m density so MLDs likely shallower.
^MLD based on sigma-t but largely driven by temperature change - if based solely on salinity then MLD is 18 m (mean T = 29.93°C).

In addition to stations where depth profiles of parameters were obtained, we report surface data for DNA samples (*Section 3.6*) from transects in the region from both the ship’s underway seawater system (∼5-m intake) and a towed surface fish starting on 26 Jan 2022 and ending on 5 March 2022 (∼1 m; Fig. 1, Supp. Table S2).

### 3.2. Phytoplankton carbon biomass

Depth profiles of phytoplankton biomass, divided into PRO, SYN and PEUK, show that PRO had the most biomass of any group in all cycles, with slightly more carbon biomass in the upper EZ (Fig. 2). SYN were a minor part of total autotrophic biomass, and autotrophic eukaryotes had ∼4-times more biomass in the lower part of the EZ, with a maximum near the DCM.

**Figure 2.**
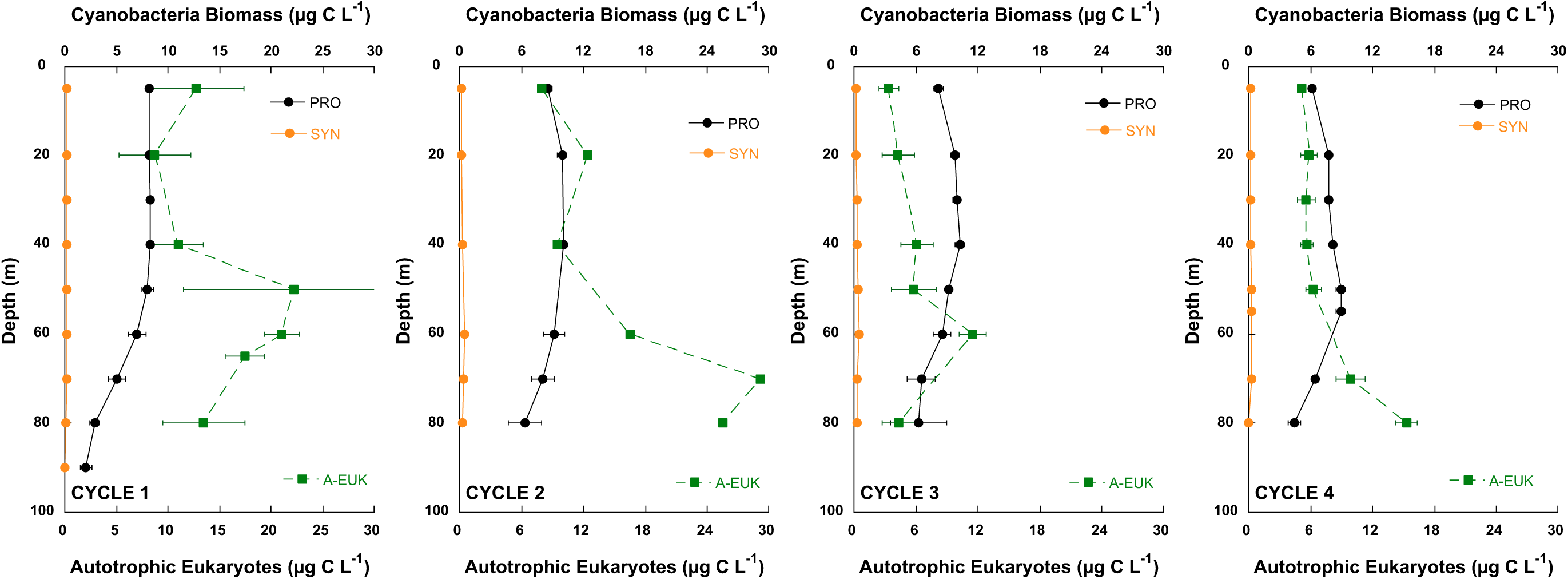
Phytoplankton biomass in each cycle, divided into cyanobacteria *Prochlorococcus* (PRO), *Synechococcus* (SYN) and autotrophic eukaryotes (A-EUK). Data are from combined flow cytometry and epifluorescence microscopy determinations.

Multiple microscopy methods (EPI-MICRO, INVERT-MICRO and IFCB) were used to determine diatom biomass, and EPI-MICRO only detected diatoms in a few samples, while INVERT-MICRO showed that the upper EZ had a diatom biomass of 0.015 ± 0.004 µg C L^-1^, and ∼5-times more biomass in lower EZ (0.071 ± 0.029 µg C L^-1^, data not shown). Surface samples (∼5-m) from the IFCB showed a much higher diatom biomass (0.88 ± 0.18 µg C L^-1^), but this was still only ∼6% of total mixed-layer autotrophs as assessed by EPI-MICRO samples.

Dinoflagellates were also determined using microscopy, however the INVERT-MICRO method does not distinguish between autotrophic and heterotrophic dinoflagellates, and the IFCB method only includes chlorophyll-containing cells. Thus, to use the INVERT-MICRO data, we used the estimate for A-DINO from EPI-MICRO (3.5 ± 0.3 µg L^-1^, assuming most micro-sized cells were dinoflagellates) and the ratio of A-DINO to total dinoflagellates (67%), to get an estimate of 1.9 ± 0.5 µg C L^-1^ for A-DINO. However, as for diatoms, the IFCB estimate was ∼10-times higher for A-DINO (21.29 ± 1.01 µg C L^-1^).

Phytoplankton carbon was integrated by depth in the EZ at each station (Table 3). Total A-EUK biomass ranged from 776-1604 mg C m^-2^, with a mean of 1229 ± 4 mg C m^-2^. PRO was the lowest percent of total autotrophic biomass in Cycle 1, at 38%, increasing to 57%, 62%, and 71% in Cycles 2, 3, and 4, respectively. MICRO-EUKS showed the opposite trend, with the highest percentage in Cycle 1 (43%) decreasing to 22%, 11% and 3% in Cycles 2, 3, and 4, respectively. In contrast, NANO-EUKS, PICO-EUKS, and SYN remained at relatively constant percentages of total biomass in all cycles, ranging from 15-21%, 3-6%, and 1-2%, respectively.

**Table 3.** Euphotic zone integrated phytoplankton biomass (mg C m^-2^) in each cycle. Depth integrals are for the depths shown, representing the euphotic zone (to 1% incident irradiance). Phytoplankton populations are *Prochlorococcus* (PRO), *Synechococcu*s (SYN), pico-eukaryotes (PICO-EUKS), nano-eukaryotes (NANO-EUKS), and micro-eukaryotes (MICRO-EUKS). PRO, SYN and PICO-EUKS are from flow cytometry data, while NANO-EUKS and MICRO-EUKS are from epifluorescence microscopy data. ND: No data available.

| Cycle | CTD | Depth interval (m) | PRO | SYN | PICO-EUKS | NANO-EUKS | MICRO-EUKS | TOTAL |
| --- | --- | --- | --- | --- | --- | --- | --- | --- |
| 1 | 11 | 0-90 | 626 | 14 | 70 | 366 | 527 | 1604 |
|  | 18 | 0-80 | 542 | 12 | 52 | 181 | 662 | 1449 |
|  | 25 | 0-80 | 597 | 11 | 38 | 180 | 729 | 1555 |
|  | 33 | 0-65 | 480 | 8 | 26 | 161 | 577 | 1252 |
|  | 38 | 0-80 | 664 | 13 | ND | ND | ND | ND |
| 2 | 48 | 0-80 | 697 | 24 | 40 | 185 | 299 | 1245 |
|  | 54 | 0-80 | 781 | 30 | 52 | 176 | 315 | 1354 |
|  | 61 | 0-80 | 818 | 26 | 27 | 226 | 185 | 1282 |
|  | 68 | 0-80 | 778 | 26 | 77 | 245 | 414 | 1541 |
|  | 74 | 0-80 | 590 | 16 | ND | ND | ND | ND |
| 3 | 86 | 0-80 | 805 | 29 | 68 | 304 | 289 | 1497 |
|  | 93 | 0-70 | 625 | 19 | 43 | 272 | 127 | 1086 |
|  | 99 | 0-60 | 592 | 18 | 37 | 142 | 15 | 804 |
|  | 106 | 0-80 | 616 | 18 | ND | ND | ND | ND |
| 4 | 114 | 0-80 | 593 | 21 | 66 | 274 | 22 | 977 |
|  | 121 | 0-80 | 596 | 17 | 41 | 113 | 10 | 776 |
|  | 128 | 0-80 | 589 | 18 | 37 | 97 | 43 | 783 |

*\*only integrated to 60 m, as deepest depth missing*

3.3. *Phytoplankton pigment biomass*

Depth profiles of DV-CHL*a* and MV-CHL*a* (the latter associated with SYN and autotrophic eukaryotes) had lower values in the upper EZ, increasing towards the lower EZ in all cycles (Fig. 3). Also, similar concentrations of DV-CHL*a* and MV-CHL*a* associated with eukaryotes were found at depths shallower than 40-50 m, with MV-CHL*a* increasing over DV-CHL*a* towards the DCM (Fig. 3). Thus, pigment results were divided into upper (<50 m) and lower (≥50 m) parts of the EZ, as these depth horizons also appeared to have different taxonomic characteristics based on accessory pigments. Depth profiles of all pigments determined by HPLC are found for each cycle in Supp. Tables S3-S6.

**Figure 3.**
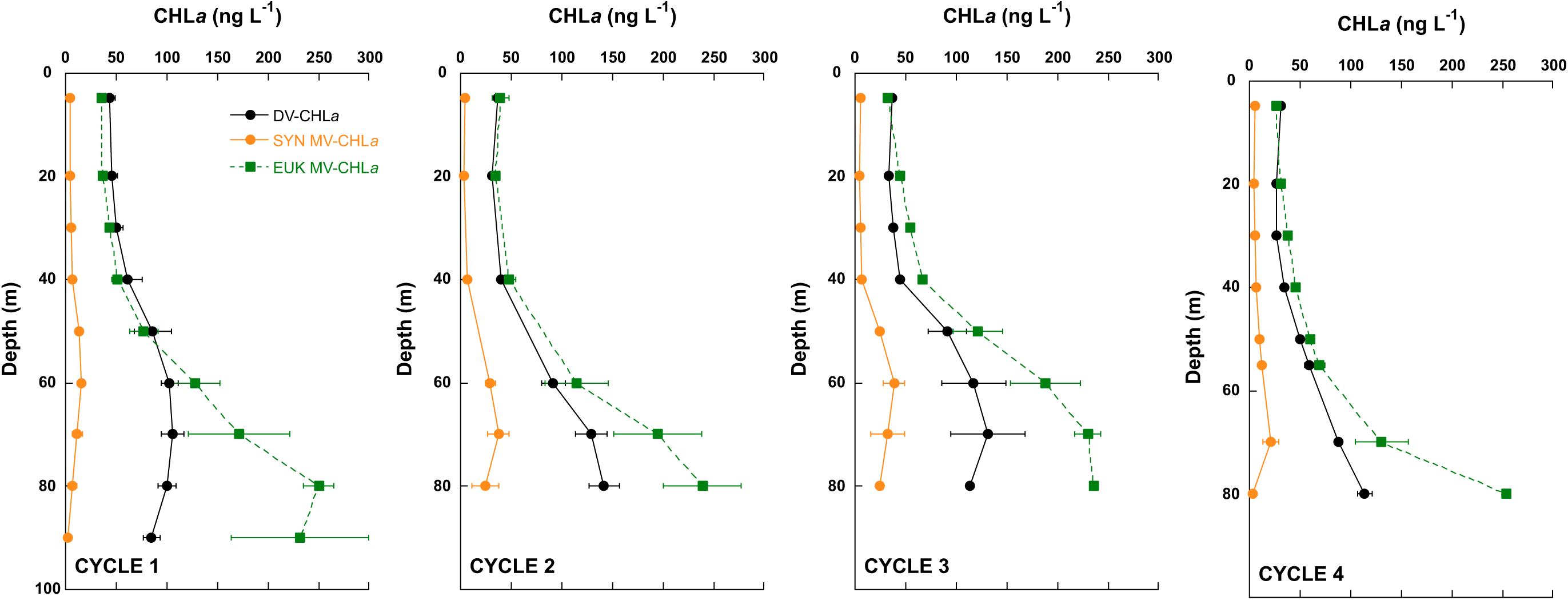
Depth profiles of chlorophyll (CHL*a*) from each cycle divided into 3 groups – divinyl chlorophyll a (DV-CHL*a*), monovinyl chlorophyll a (MV-CHL*a*) associated with *Synechococcus* (SYN MV-CHL*a*), and MV-CHL*a* associated with all other eukaryotic phytoplankton (EUK MV-CHL*a*).

The signature pigment of PRO, DV-CHL*a*, was a large portion of *T*CHL*a* (46% in the upper EZ and 35% in the lower EZ) and ranged from ∼29 to 54 ng L^-1^ in the upper EZ (Table 4). DV-CHL*a* was much higher in the lower EZ at ∼86 to 132 ng L^-1^ (Table 4). MV-CHL*a*, which is found in all other groups of phytoplankton, was also lower in the upper EZ (∼39 to 54 ng L^-1^) than in the lower EZ (159 to 240 ng L^-1^).

**Table 4.**
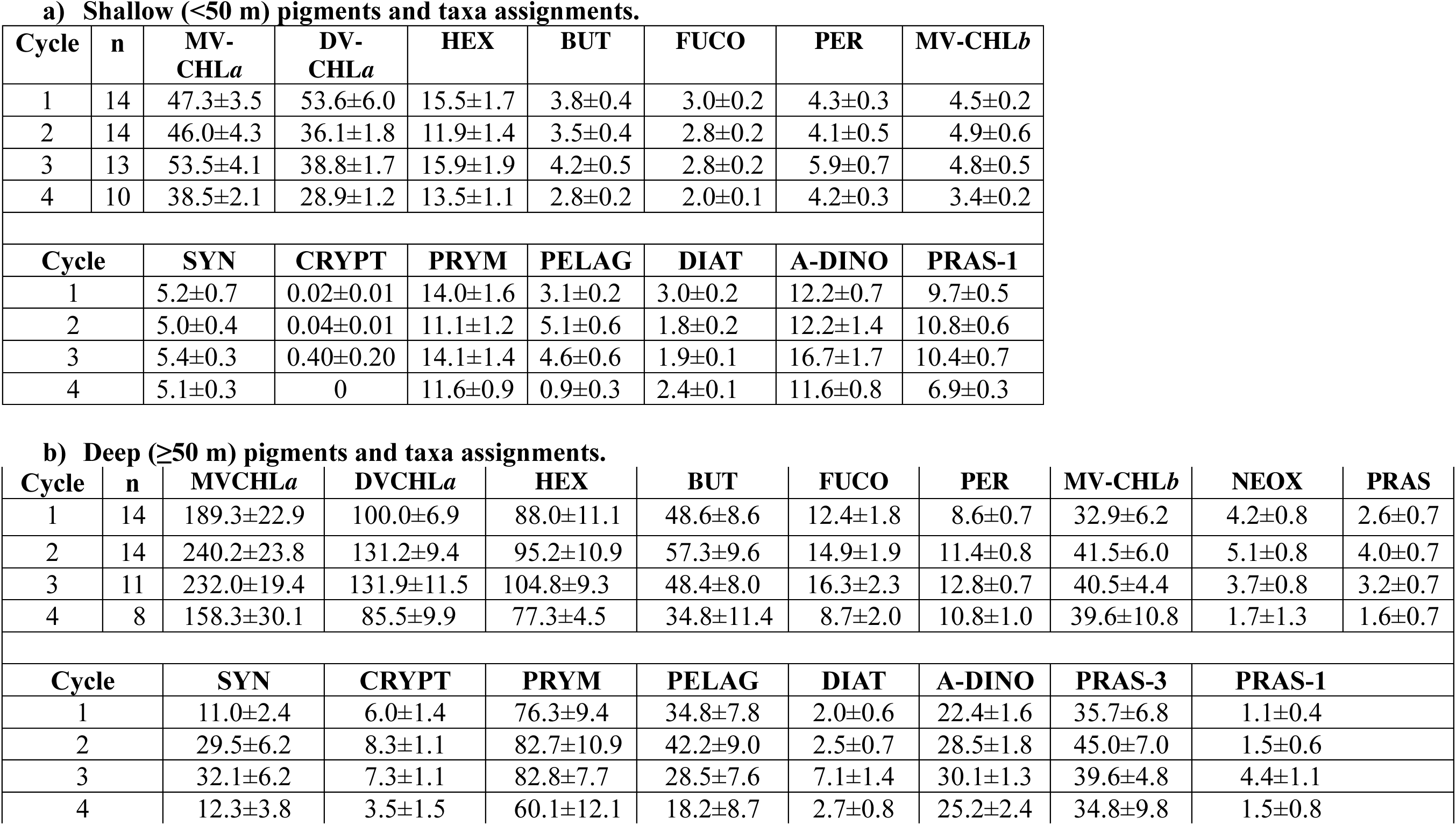
a) Shallow (<50 m) and deep (≥50 m) pigments (ng L^-1^) and taxa (ng MV-CHLa L^-1^) assignments. Pigments are from HPLC determinations and taxa assignments are from phytoclass (ver. 1.0.0). Data are averages ± 1 standard error. No prasinoxanthin was detected in shallow samples, and only pigments that were >1 ng L^-1^ are shown, however allophycocyanin, neoxanthin, violaxanthin, and lutien (all <1 ng L^-1^) were used to help determine taxonomic assignments.

Accessory pigments were used to partition MV-CHL*a* into different taxonomic groups and, as for MV- and DV-CHL*a*, these pigments were much higher in the lower EZ. In the upper EZ, the main taxonomic groups present, in order of their share of MV-CHL*a*, were PRYM, A-DINO, and PRAS-1, followed by SYN, PELAG, DIAT, and CRYPT (Table 4). In the lower EZ, PRYM was again a prominent group, but PELAG and PRAS-3 were more important than A-DINO, followed by SYN, CRYPT, DIAT, with PRAS-1 representing the lowest amount of MV-CHL*a* (Table 4).

Depth profiles of all taxonomic assignments for each cycle are found in Supp. Tables S7-S10. PRYM and PELAG taxa were mostly associated with the accessory pigments HEX and BUT, respectively; however, these two taxonomic groups also each have the other pigment in their makeup as well as, such as FUCO (Supp. Table S2). Nevertheless, in all cycles and at all depths, HEX and BUT pigment concentrations paralleled their PRYM and PELAG taxonomic assignments to MV-CHL*a* (Fig. 4), with PRYM and PELAG averaging 31 ± 1% and 9 ± 1% of MV-CHL*a*, respectively (Fig. 4, Supp. Table S7-S10). Depth profiles of PRAS (belonging to the chlorophyte group) were divided into the PRAS-3 taxon with the pigment PRAS and PRAS-1 taxa which do not have PRAS but do have MV-CHL*b* (Fig. 5). PRAS-3 was absent in the upper EZ, increasing towards the DCM (lower EZ average of 16.2 ± 0.1% of MV-CHL*a*), whereas PRAS-1 had the opposite distribution – high in the upper EZ (20 ± 0.4% of MV-CHL*a*) and nearly absent in the lower EZ. A-DINO were assigned to MV-CHL*a* based on their PER content (Supp. Table S2) and tracked that pigment well (Fig. 6). At 5 m, A-DINO were 26 ± 1% of MV-CHL*a*, whereas they averaged 23 ± 1% for all depths (Supp. Table S7-S10). DIAT and CRYPT represented a relatively low amount of the total MV-CHL*a*, and both taxa increased towards the DCM (Fig. 7). CRYPT, whose indicator pigment is ALLO, tracked that pigment well (Fig. 7). In contrast, FUCO, the indicator pigment for diatoms, did not parallel the MV-CHL*a* assigned to them well – potentially because FUCO is also found in the more dominant taxa PRYM and PELAG. For comparison, the 5-m average amount of MV-CHL*a* assigned to diatoms was 3 ± 0.2%, which is about half of the IFCB estimated contribution to carbon biomass for this group.

**Figure 4.**
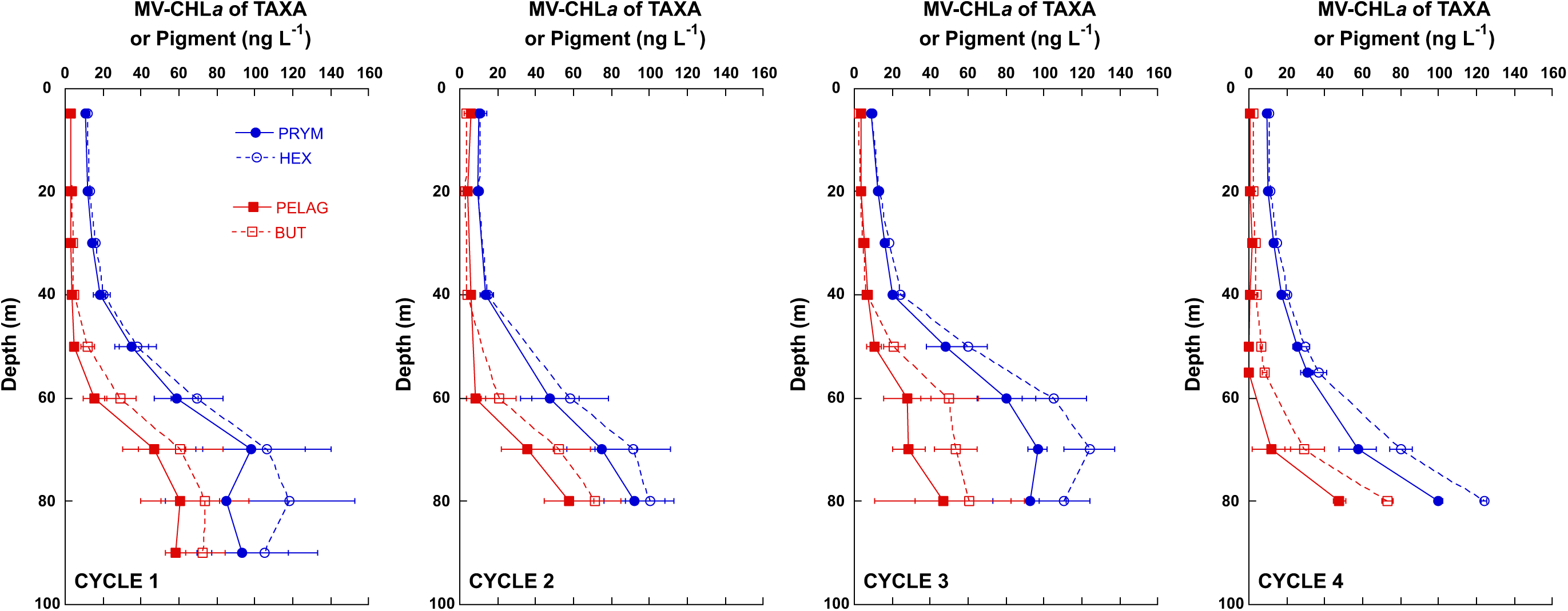
Depth distributions in each cycle of the monovinyl chlorophyll a (MV-CHL*a*) associated with prymnesiophytes (PRYM) and pelagophytes (PELAG), and their indicator pigments 19’-hexanoyloxyfucoxanthin (HEX) and 19’-butanoyloxyfucoxanthin (BUT).

**Figure 5.**
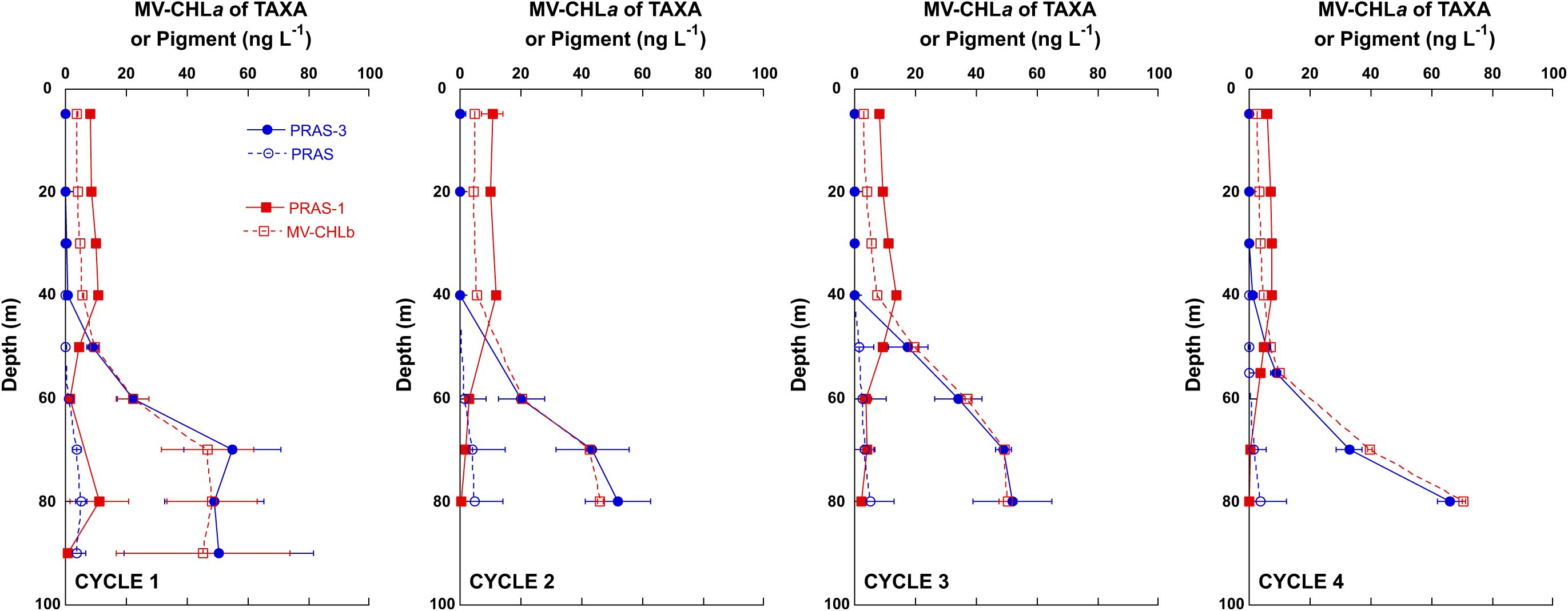
Depth distributions of the monovinyl chlorophyll *a* (MV-CHL*a*) associated with prasinophytes and their indicator pigments prasinoxanthin (PRAS) and monovinyl chlorophyll *b* (MV-CHL*b*) in each cycle. Prasinophyte Group 3 (PRAS-3) is associated with PRAS, and prasinophyte group 1 (PRAS-1) is associated with MV-CHL*b*.

**Figure 6.**
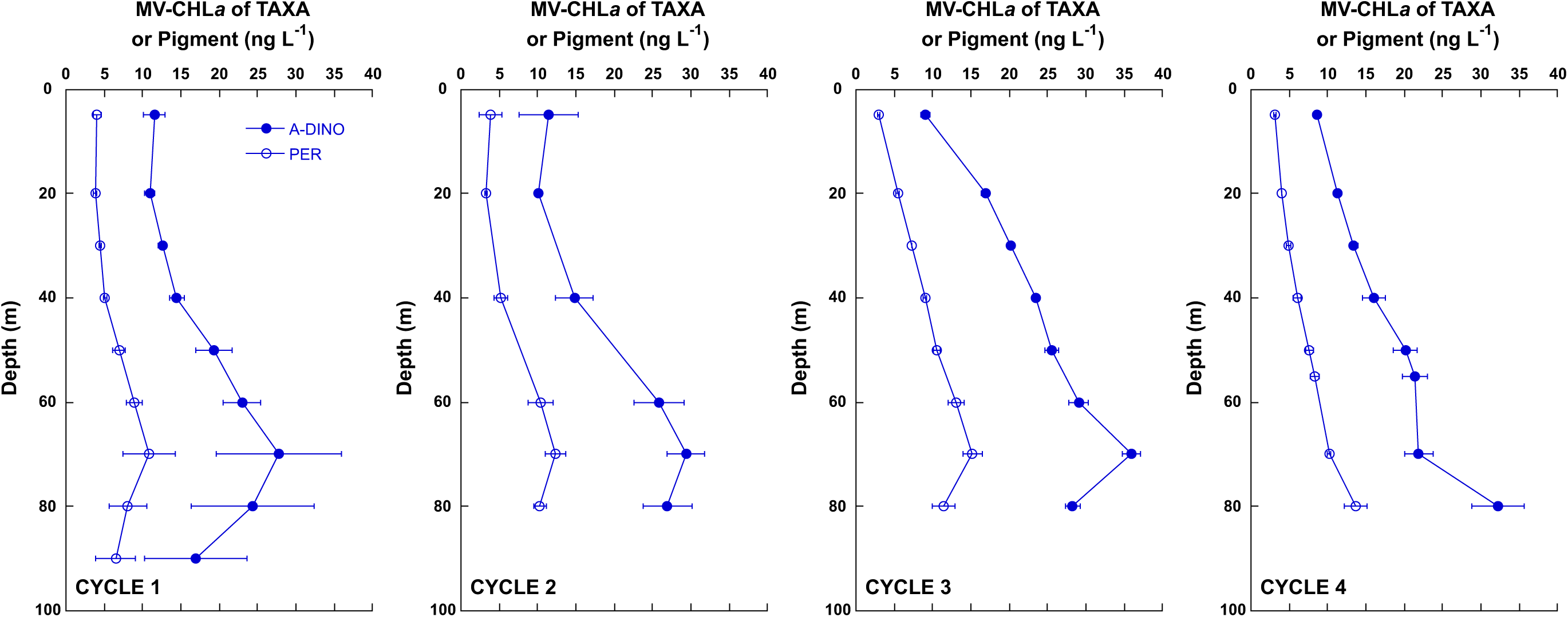
Depth profiles from each cycle of the MV-CHL*a* associated with autotrophic dinoflagellates (A-DINO) and their indicator pigment peridinin (PER).

**Figure 7.**
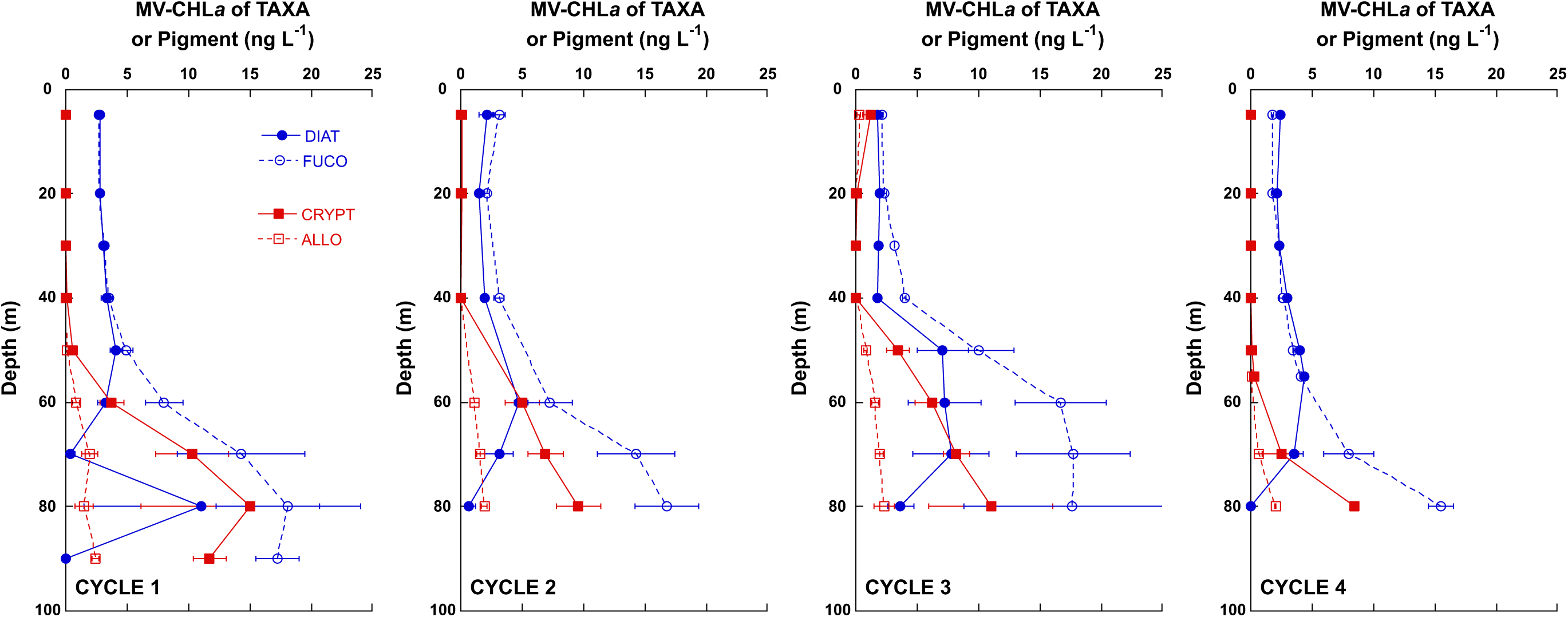
Depth profiles from each cycle of the monovinyl chlorophyll a (MV-CHL*a*) associated with diatoms (DIAT) and cryptophytes (CRYPT), and their indicator pigments fucoxanthin (FUCO) and allophycocyanin (ALLO).

All pigments were integrated by depth over the EZ (Table 5). Overall, mean *T*CHL*a* was 12 ± 0.7 µg m^-2^, with MV-CHL*a* and DV-CHL*a* comprising 61% and 39% of that mean, respectively. HEX was 25% of *T*CHL*a*, with BUT and MV-CHL*b* ∼10% each. FUCO, PER and PRAS were ≤4% of *T*CHL*a*. No particular trend in pigment concentrations was observed; however, Cycle 4 values were consistently lower than other cycles.

**Table 5.** Euphotic zone integrated pigment biomass (ng m^-2^). Pigments are from discrete HPLC samples taken over the depth range indicated (m). Pigment abbreviations are defined in Table 1.

| Cycle | CTD | Depth interval | TCHLa | MV-CHLa | DV-CHLa | HEX | BUT | FUCO | PER | MV-CHLb | PRAS |
| --- | --- | --- | --- | --- | --- | --- | --- | --- | --- | --- | --- |
| 1 | 11 | 0-90 | 15373 | 9887 | 5482 | 4326 | 2374 | 699 | 548 | 1869 | 135 |
|  | 18 | 0-80 | 12529 | 8838 | 4590 | 3733 | 1935 | 590 | 504 | 1192 | 98 |
|  | 25* | 0-60 | 7961 | 3554 | 4409 | 1383 | 329 | 222 | 311 | 345 | 0 |
|  | 33 | 0-65 | 10023 | 5356 | 4667 | 2423 | 945 | 291 | 362 | 645 | 24 |
|  | 38 | 0-90 | 14426 | 8326 | 6096 | 3454 | 1520 | 474 | 486 | 1547 | 86 |
| 2 | 48 | 0-80 | 14801 | 8623 | 6173 | 2827 | 1383 | 542 | 593 | 1237 | 60 |
|  | 54 | 0-80 | 11291 | 6443 | 4846 | 1919 | 766 | 353 | 488 | 867 | 85 |
|  | 61 | 0-90 | 13554 | 7979 | 5580 | 2669 | 1221 | 465 | 556 | 1147 | 136 |
|  | 74 | 0-80 | 16867 | 11266 | 5604 | 4688 | 2449 | 658 | 620 | 1667 | 152 |
| 3 | 86 | 0-80 | 16602 | 10065 | 6542 | 3941 | 1489 | 687 | 698 | 1390 | 98 |
|  | 93 | 0-70 | 14509 | 8558 | 5953 | 3419 | 1305 | 599 | 586 | 1274 | 90 |
|  | 99 | 0-60 | 8051 | 4947 | 3103 | 1903 | 762 | 281 | 414 | 611 | 18 |
|  | 106 | 0-80 | 13612 | 8972 | 4640 | 3836 | 1649 | 577 | 763 | 1370 | 76 |
| 4 | 114 | 0-80 | 9896 | 5900 | 3996 | 2613 | 822 | 301 | 521 | 1313 | 28 |
|  | 121 | 0-80 | 10812 | 6848 | 3965 | 3080 | 1282 | 398 | 501 | 1049 | 60 |
|  | 128 | 0-80 | 10089 | 6142 | 3949 | 2811 | 873 | 307 | 581 | 1117 | 23 |

\*only integrated to 60 m, as deepest depth missing

Summarizing the contributions of the different taxa to the entire phytoplankton community for all cycles as proportions of *T*CHL*a* in the upper (shallow) EZ, PRO was the dominant group, followed by PRYM, A-DINO, and PRAS-1, with more minor contributions of SYN, PELAG, and DIAT (Fig. 8a). In the lower EZ, PRO, PRYM, and A-DINO were still the main taxa, but PRAS-3 replaced PRAS-1 as a dominant, and a higher proportion of MV-CHL*a* was attributed to PELAG, CRYPT, and DIAT (Fig. 8b).

**Figure 8.**
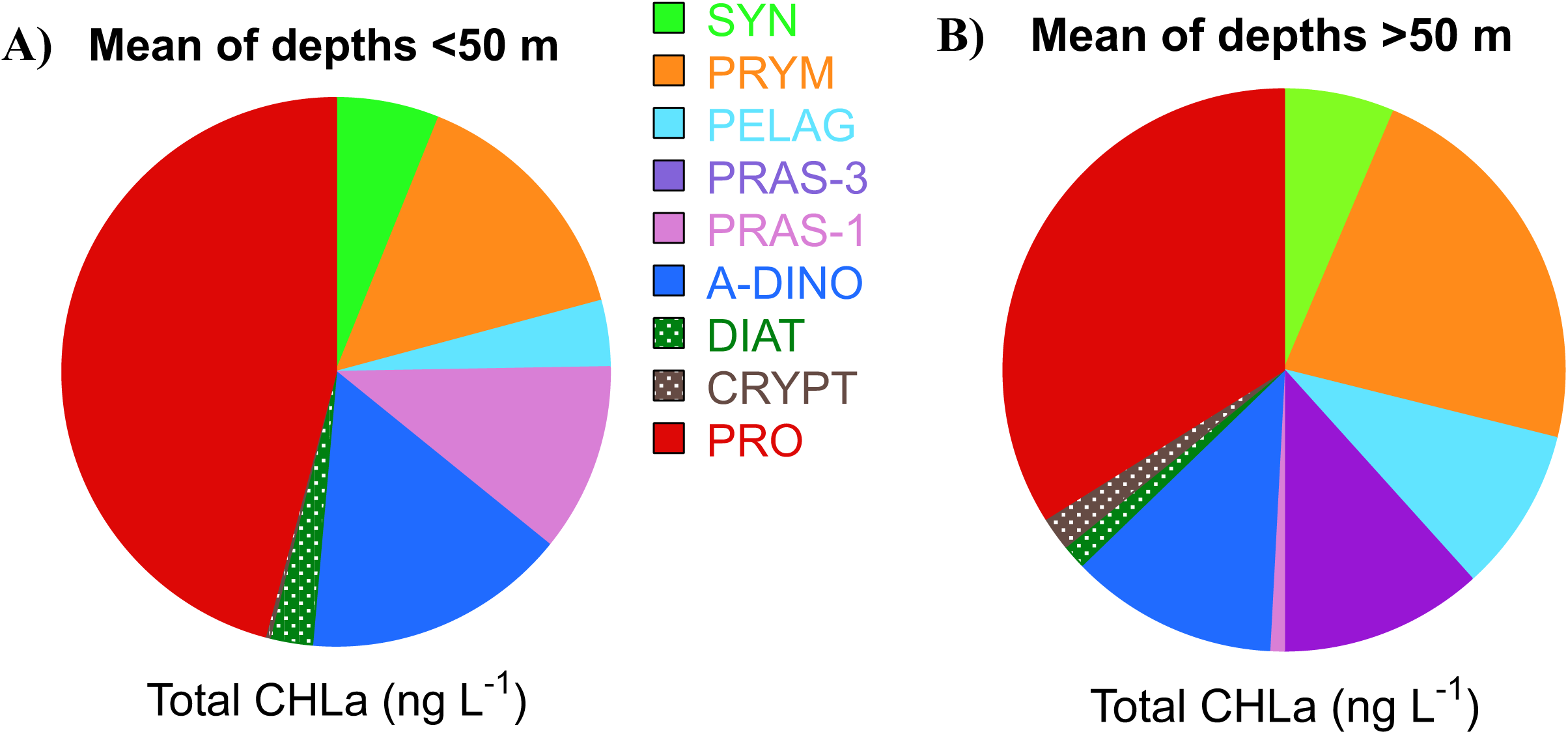
Pie charts showing the mean percent composition (of total CHL*a*, *T*CHL*a*) of the phytoplankton community in two depth horizons: A) shallower (upper euphotic zone, <50 m) and B) deeper (lower euphotic zone, >50 m). Abbreviations for different taxa are defined in Table 1. Mean *T*CHL*a* (± 1 standard error) of all cycles of the shallow and deep communities are 86 ± 7 ng CHL*a* L^-1^ and 331 ± 40 ng CHL*a* L^-1^, respectively.

### 3.4. Phytoplankton carbon:CHLa ratios

Using the carbon and pigment biomass estimates described above, we estimated C:CHL*a* ratios for each depth profile, and averaged all values for upper and lower EZ depths (Table 6). The C:*T*CHL*a* ratios for the entire community were 120 ± 8, 173 ± 8, and 60 ± 6 for the entire, upper and lower EZ depths, respectively. Dividing *T*CHL*a* into its DV- and MV-CHL*a* components revealed that PRO had the highest C:DV-CHL*a* value of 162 ± 11 for all depths in the EZ, 237 ± 10 in the upper EZ, dropping to 74 ± 8 in the lower EZ. The C:MV-CHL*a* ratios, which mostly represent eukaryotic phytoplankton since SYN biomass was generally low, were much lower at 90 ± 9, 126 ± 13, and 51 ± 8 for the entire, upper and lower EZs, respectively.

**Table 6.** Euphotic zone (to 1% light level) Carbon:Chlorophyll *a* (C:CHL*a*) ratios. Shown are mean C:CHL*a* ratios of overall (all euphotic zone depths), shallow (upper euphotic zone, <50 m) and deep (lower euphotic zone, ≥50 m) mean values. For the ratios: total autotrophic carbon is relative to total chlorophyll *a* (*T*CHL*a*), PRO carbon is relative to divinyl chlorophyll *a* (DV-CHL*a*), and eukaryotic + SYN carbon is relative to monovinyl chlorophyll *a* (MV-CHL*a*). N = number of determinations; errors are ±1 standard error of the mean.

| EUPHOTIC ZONE DEPTHS | N | C:TCHL*a* | C:DV-CHL*a* | C:MV-CHL*a* |
| --- | --- | --- | --- | --- |
| Overall | 80 | 120 ± 8 | 162 ± 11 | 90 ± 9 |
| Shallow (<50 m) | 42 | 173 ± 8 | 237 ± 10 | 126 ± 13 |
| Deep (≥50 m) | 38 | 60 ± 6 | 74 ± 8 | 51 ± 8 |

### 3.5. Phytoplankton distributions: mixotroph abundance

Most eukaryotic taxonomic groups are capable of both photosynthesis and phagotrophy, with the exception of diatoms. Thus, many of the eukaryotic taxa described above will have mixotrophic members (MEUK), and a subset of our samples were analyzed for the presence of acidic vacuoles that would indicate active phagotrophy (Selph et al., this issue).

In Cycles 1-4, MEUK were a higher percentage of chlorophyll-bearing eukaryotic cell abundance at depths shallower than 55 m (upper EZ, means of 35 to 84%), relative to deeper samples (lower EZ, 60-80 m) where they ranged from 30 to 51% (Table 7). The percent of MEUK increased temporally, with higher abundances found in Cycles 3 and 4 compared to earlier cycles. These later cycles were characterized by stronger stratification and warmer surface temperatures.

**Table 7.** Cycle averages of mixotrophic eukaryote (MEUK) abundance as a percent of total chlorophyll-bearing eukaryotic cells (MEUK + photosynthetic eukaryotes) in shallow depths (<55 m) and deep depths (≥60-80 m) of the euphotic zone. MEUK abundances were determined using flow cytometry of living cells stained with LysoTracker Green (Selph et al., this issue, a). Errors are ± 1 standard error of the mean.

| **CYCLE** | **Shallow ($\leq 55$ m)** | **Deep ($\geq 60$-80 m)** |
| --- | --- | --- |
| 1 | $48 \pm 2\%$ | $30 \pm 3\%$ |
| 2 | $35 \pm 2\%$ | $30 \pm 4\%$ |
| 3 | $58 \pm 5\%$ | $45 \pm 5\%$ |
| 4 | $84 \pm 1\%$ | $51 \pm 6\%$ |

### 3.6. Phytoplankton DNA-based abundances and diversity

Absolute abundances of both 16S and 18S rRNA gene copies were estimated to examine the abundance and diversity of the prokaryotic (cyanobacterial) and eukaryotic phytoplankton communities within the mixed layer and DCM (Fig. 9). In line with other measurements, PRO dominated the cyanobacteria, averaging greater than 98.6% (3.34 – 7.21 x 10^9^) of the copies L^-1^ within each cycle or transect station. PRO was also the most dominant group in the DCM; however, 11% and 23% of the cyanobacterial community in Cycles 1 and 2, respectively, were unclassified *Synechococcales*, which includes both PRO and SYN. Within eukaryotes, A-DINO dominated the community with average abundances during the cycles ranging from 1.24 x 10^8^ to 1.70 x 10^8^ copies L^-1^, except during Cycle 3 where average abundances were highest at 1.79 x 10^8^ and 2.61 x 10^8^ copies L^-1^ in the mixed layer and DCM, respectively. However, 66% of A-DINO ASVs, representing 51% of the total A-DINO 18S copies, were not annotatable to the genus level.

**Figure 9.**
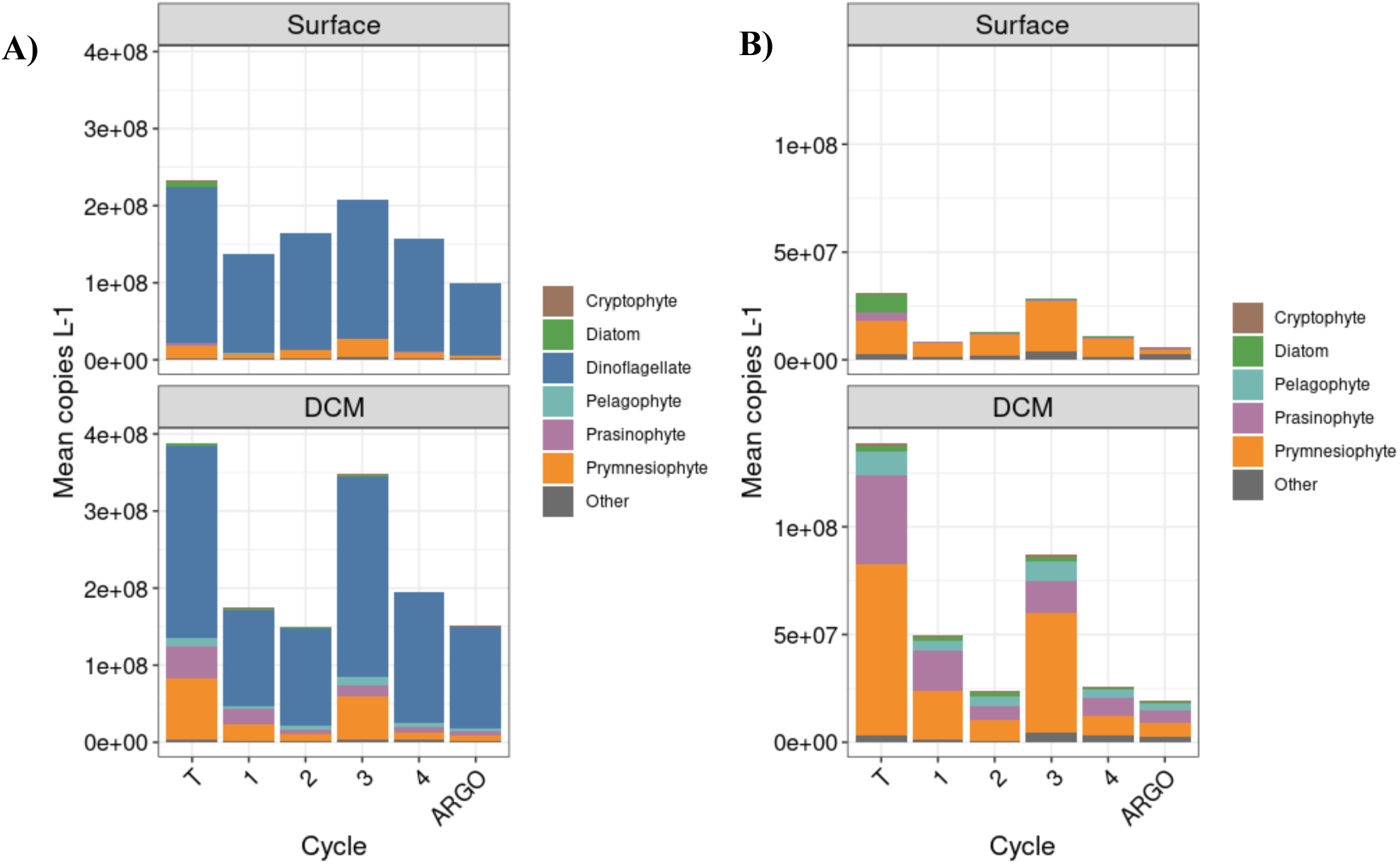
Mean copies of 18S rRNA gene (copies L^-1^) in each group for each cycle, Argo stations, or transect samples within the mixed layer or DCM. A) All population groups, B) All population groups except dinoflagellates.

Corresponding to high average A-DINO abundances in Cycle 3, the highest community-wide abundances were also observed during Cycle 3 (Fig. 9). In the surface samples, all groups increased in abundance except DIAT, which had the highest abundances during Cycle 4. In the DCM, all groups had the highest abundances during Cycle 3, except prasinophytes and DIAT, which were both highest during Cycle 1. In comparing mean group abundances between the surface and DCM, prasinophytes, CRYPT, DIAT, and PELAG were consistently higher in the DCM. PRYM abundances were higher in the DCM during Cycle 1 and 3, but higher in the surface for Cycles 2 and 4. Within DIAT, pennate taxa comprised 50% and 58% of reads during Cycles 1 and 2, respectively, and increased to 75% and 72% during Cycles 3 and 4, respectively.

Alpha diversity of the eukaryotic phytoplankton community, expressed as the Shannon index, was not significantly different among cycles and depth horizons (Supp. Fig. S2A, P > 0.05, Kruskal-Wallis test). However, the surface communities were significantly different from the DCM communities (Supp. Fig. S2B, Bray-Curtis, PERMANOVA, P < 0.05). Most phytoplankton genera were shared among depth horizons; however, several rarer genera were specific to each. Dominant genera in both depth horizons included the A-DINO *Gyrodinium*, *Gymnodinium, Prorocentrum, Warnowia, Karlodinium, Lepidodinium*; the PRYM *Chrysochromulina* and *Phaeocystis*, and the prasinophyte genus *Ostreococcus*. Surface communities had high abundances of the A-DINO genera *Tripos* and *Margalefidinium*, as well as the DIAT genus *Pseudo-nitzschia*, whereas DCM communities had high abundances of the PELAG species *Pelagomonas caleolata*, the bolidophyte order *Parmales*, and the prasinophyte genus *Bathycoccus*, in addition to the aforementioned high abundance of *Ostreococcus*.

While the DCM communities were not significantly different from one another, the surface communities in Cycle 1 were significantly different from all other cycles (P < 0.05, Supp. Fig. S2B). Additionally, the surface community of Cycle 2 was distinct from that of Cycle 3 (P = 0.017). In Cycle 1, unclassified *Gymnodiniales* and the genus *Warnowia* within A-DINO had approximately 2-8x greater relative abundances compared to other cycles (Supp. Fig. S3). When examining the genus-level where possible, Cycle 2 was largely similar to Cycle 3, suggesting differences on the species or strain level (Supp. Fig. S3).

Both the 16S and 18S rRNA gene data were investigated to examine the presence of either photoautotrophic diazotrophs or eukaryotic phytoplankton that may have symbiotic diazotrophs. *Trichodesmium* was detected during all cycles, but was normally less than 1% of the cyanobacterial reads. Within two transect samples, *Trichodesmium* represented 7% and 26% of cyanobacterial reads, suggesting that there were infrequent patches of increased *Trichodesmium* abundances. During cycles, the other cyanobacterial diazotrophs *Richelia, Crocosphaera, Epithemia* endosymbionts, and UCYN-A were detected in low abundances (<1% of cyanobacteria) in only 5 or fewer samples. *Calothrix* reads were undetected.

Members of the diatom genera *Hemiaulus* and *Rhizosolenia* may contain *Richelia* as endosymbionts at times (Pierella Karlusich et al., 2021). *Rhizosolenia* was detected in Cycles 1-3, and microscopy samples showed *Richelia* endosymbionts. Both *Hemiaulus* and *Epithemia*, which have a different nitrogen-fixing endosymbiont, were only detected during some transect samples. Moreover, there was little overlap between samples where these diatom genera were detected and their respective endosymbionts. *Braarudosphaera*, which may contain nitroplasts or UCYN-A (Coale et al., 2024), were each detected in 7 mixed layer samples during the transects and Cycles 2 and 3, but only with some overlap.

## 4. Discussion

### 4.1. Study overview

The Argo Basin, a 5000-m deep area with low CHL*a* and nutrients, is downstream of the Indonesian Throughflow. The phytoplankton community in this area is little studied, making this report its first comprehensive description. We used multiple independent, but complementary, methods to determine phytoplankton community composition. Samples were collected for taxonomic identification using DNA, pigments with HPLC, and 2 flow cytometric methods - one for determining abundances of pico and nano-plankton and one for assessing the presence of acidic vacuoles for phagocytosis. To determine the abundance and biomass of the larger (micro) members of the microbial community, 3 microscopic approaches were used – 1) epifluorescence microscopy for nano- and micro-plankton, 2) inverted microscopy for diatoms and dinoflagellates, and 3) an underway surface sampler (Imaging FlowCytobot) to describe surface populations of >8-µm chlorophyll-bearing cells.

The data presented here was gathered during the austral summer, ∼1 week after a tropical storm went through our first cycle’s station area. Subsequently the water column stabilized with mixed layers shoaling, while the surface showed gradually increasing temperatures up to 30.55°C. We found that this region was characterized by a phytoplankton community typical of warm, nitrogen-limited conditions, and in the following discussion compare our results to similar environments in the GoM and the subtropical Pacific HOT site.

### 4.2. Overall phytoplankton community

The composition of the phytoplankton community found in the Argo Basin is similar to that in other open ocean oligotrophic habitats, such as the HOT site in the Pacific Ocean (Andersen et al., 1996), the oligotrophic sites visited during the Malaspina expedition (Latasa et al., 2023), and the offshore land-remote deeper waters of the Gulf of Mexico (Selph et al., 2021). In all of these locations, PRO is the numerical dominant, and pico- and nano-sized cells comprise the majority of the eukaryotic community, with only small contributions of larger micro-sized diatoms. As indicated in Kranz et al. (this issue), primary productivity across the cycles remained robust, with depth-integrated net primary productivity (NPP) around 460 mg C m^-2^ d^-1^, comparable to other oligotrophic ocean sites such as HOT and the Bermuda Atlantic Times-series Study (Brix et al., 2006).

Mean upper-EZ *T*CHL*a* in the Argo Basin was 0.085 µg L^-1^, compared to ∼0.075 µg L^-1^ at the HOT site in summer (Pasulka et al., 2013), and 0.038 µg L^-1^ at the GoM site (Selph et al., 2021). In the lower EZ, *T*CHL*a* averaged 0.32, ∼0.31, and 0.27 µg L^-1^ for the Argo Basin, HOT, and GoM, respectively. Despite the Argo Basin having 30% higher average *T*CHL*a* values, its integrated EZ *T*CHL*a* was ∼13 mg m^-2^, similar to the GoM (10 mg m^-2^), but much lower than at HOT (∼30 mg m^-2^ in summer). The HOT higher integral is a result of its much deeper EZ, suggesting lower abiotic particle loads at HOT compared to the other two areas.

Higher biomass of photosynthetic eukaryotes was found in the lower EZ, and the composition of deeper community differed from the upper EZ. PRO, PRYM, and A-DINO together made up ∼75% of *T*CHL*a* in the upper EZ, but in the lower EZ decreased contributions of PRO and A-DINO combined with increased PRYM and PELAGO, and the addition of CRYPT made the community more diverse. Also, prasinophyte groups changed between upper and lower EZ depths. DIAT were low overall, but of the few cells seen with microscopy, many had nitrogen-fixing symbionts. *Trichodesmium* was largely absent from microscopy samples, but was widely detected in DNA samples, albeit in low abundances along with sporadic detection of other nitrogen-fixing organisms or potential hosts.

Studies of mixotrophic plankton in oligotrophic habitats (Sanders et al., 2000; Unrein et al., 2007; Zubkov and Tarran, 2008; Hartmann et al., 2013; Livanou et al., 2019) have shown that a large proportion of the community was represented by mixotrophs, as we found in this study. Mixotrophic organisms are postulated to become more heterotrophic at high temperatures (Wilken et al., 2013), which is also consistent with environmental conditions found later in the cruise (later cycles) with warmer surface waters, shallower mixed layers and higher percentage of cells with acidic vacuoles. Generally, though, this strategy is thought to augment nutrition under low-nutrient conditions. Mixotrophs were more abundant in the upper EZ relative to the lower EZ, consistent with consumption of bacteria or PRO where nitrate is low. Even at the DCM, there was a high percent of mixotrophs, and the N:P ratio was still lower than the Redfield ratio (Kranz et al., this issue), suggesting continued N-limitation at that depth. In the following sections, we discuss phytoplankton taxonomic groups in more detail.

### 4.3. Cyanobacteria

PRO dominated biomass, likely reflecting its ability to outcompete other groups when nitrate is low at high light due to its use of ammonium, small cell size which provides favorable nutrient uptake kinetics, and highly streamlined genome (Partensky et al., 1999; Partensky and Garczarek, 2010). PRO biomass varied little with depth (Fig. 2), with DV-CHL*a* increasing only ∼3 fold between the upper and lower EZ, whereas MV-CHLa increased over 4-fold (Table 4). This resulted in PRO comprising a smaller proportion of *T*CHL*a* in the lower EZ (Fig. 8). The integrated cell concentration of PRO in the Argo Basin averaged 20.3 x 10^12^ cells m^-2^, which is similar to HOT and higher than in the GoM (∼22 and 10-17 x 10^12^ cells m^-2^, respectively).

SYN were present, but at a very low abundance, as found at HOT in the subtropical Pacific (Table 8). Thus, SYN was a smaller proportion of cyanobacteria (PRO + SYN) than in the GoM (Selph et al., 2021). These data suggests that the EZ was even more nutrient impoverished in the Argo Basin than the GoM. Consistent with this, SYN is known to favor meso- or even eutrophic environments (Partensky et al., 1999; Latasa et al., 2010).

**Table 8.**
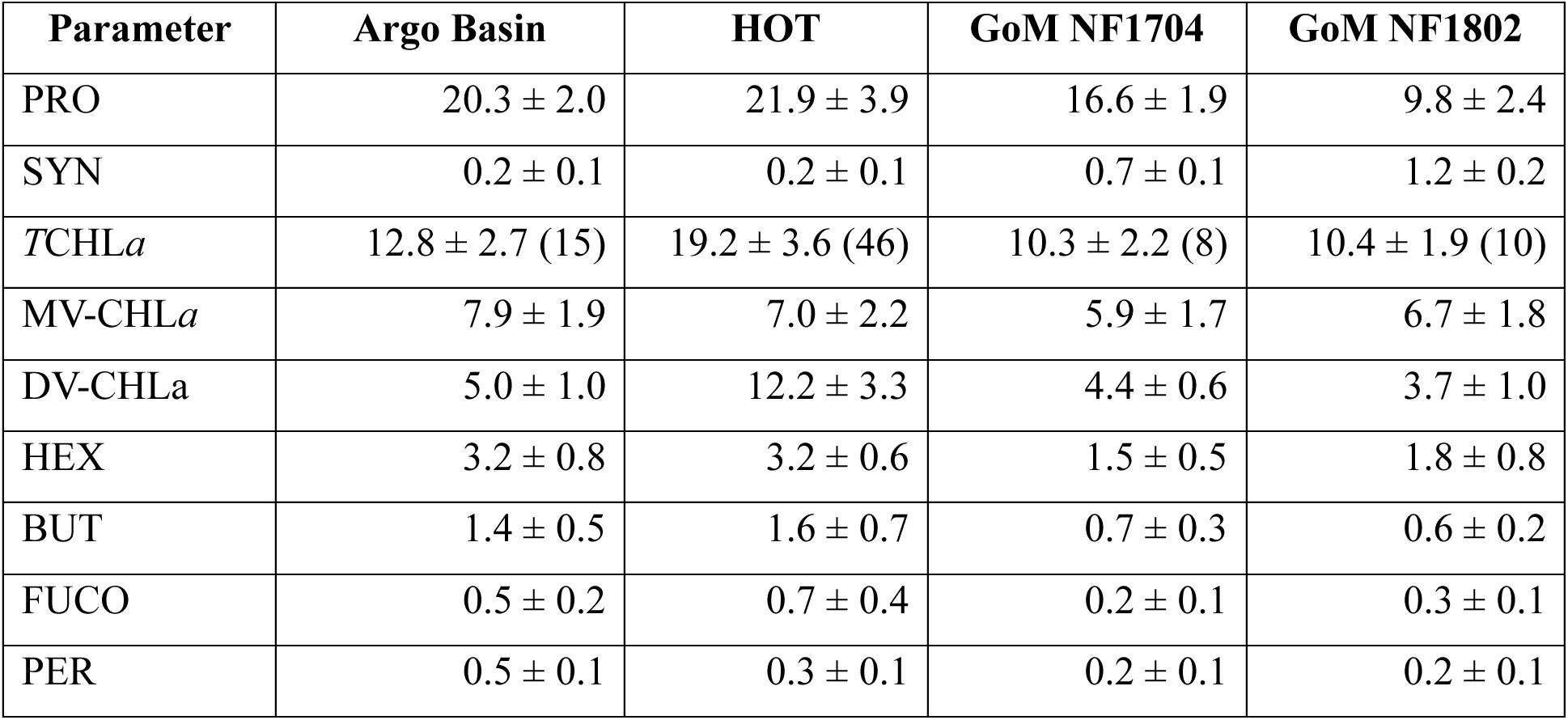
Comparisons of Argo Basin phytoplankton to the Hawaii Ocean Time-series site (HOT), and the remote, open-ocean Gulf of Mexico (GoM). All data are integrals to the 1% incident irradiance depth; for HOT this is estimated as 115 or 125 m, whichever depth was closest to the deep chlorophyll maximum depth. HOT data are from cruises 289-339 (Jan 2017 – Sept 2022, n = 46) accessed through the HOT-DOGS website; GoM data are from Table VI in Selph et al. (2021). Pigments are means based on the number of determinations indicated in parentheses for *T*CHL*a*. Uncertainties are standard deviations of the mean estimates. Pigment abbreviations are defined in Table 1.

As noted above, large quantities of *Trichodesmium* were not observed, although they did appear sporadically in surface samples throughout the cruise and were widely detected, but at low abundances, by 16S rRNA gene sequencing. We did sample specifically for them, using noon CTD casts, filtering large volumes at many depths in the upper EZ. From these samples, we saw evidence for nitrogen-fixing symbionts in several diatom species, as well as orange-fluorescing cells (with 488 nm excitation) that may have been *Crocosphaera* (data not shown).

*Crocosphaera*, UCYN-A, as well as *Richelia*, and *Epithemia* symbionts were also sporadically detected in DNA samples (data not shown). Consistent with this, nifH gene sequencing from the DNA samples used here showed nitrogen-fixing taxa were present, largely comprising *Trichodesmium* and nitrogen-fixing gammaproteobacteria (Pers. Comm., Hai Kong Io). N_2_ fixation rate measurements indicate that approximately 16% of the NPP was supported by direct or indirect N_2_ fixation (Kranz et al., this issue) and hotspots of N_2_ fixation were observed sporadically (deSouza et al., this issue).

In terms of EZ-integrated pigments, *T*CHL*a* was similar between the Argo Basin and the GoM (∼10-13 mg m^-2^), but somewhat higher at HOT (∼19 mg m^-2^), with this trend driven by DV-CHL*a* (∼4-5 mg m^-2^ in the Argo Basin and GoM, but ∼12 mg m^-2^ at HOT) with MV-CHL*a* similar between sites (∼6-8 mg m^-2^, respectively, Table 8). The lower ratio of DV-CHL*a*:PRO biomass in the Argo Basin and the NF1704 GoM cruise relative to the other sites might reflect slower growth rates of PRO (Landry et al., 2022; Landry et al., this issue) or different PRO clade dominance with lower DV-CHL*a* content per cell.

### 4.4. Eukaryotic groups

#### 4.4.1. Autotrophic dinoflagellates

The independent, complementary methods we used in this study resulted in a consistent community composition overall; however, microscopy, HPLC and DNA methods deviated for estimates of A-DINO. There was not a great increase in A-DINO in the upper EZ versus the lower EZ with the pigment data, agreeing roughly with the DNA pattern of A-DINO gene copies L^-1^. However, the DNA data showed diverse A-DINO genera being the overwhelmingly dominant group (Fig. 9A, Supp. Fig. S3), whereas HPLC data showed A-DINO comprising an average of 23% of MV-CHL*a* for all depths, which was less than the contribution of PRYM to MV-CHL*a*, especially in the lower EZ (Table 4). The high percentage of A-DINO likely contributed to the relatively high gross primary productivity:NPP (GPP:NPP) ratio (∼1.6), with high respiration rates often observed in dinoflagellates (Pitcher and Probyn, 2016; Bercel and Kranz, 2019).

Notably, eukaryotic phytoplankton taxa, comprised of many distinct groups, have widely different 18S gene copy numbers. Thus, abundant genes might reflect numerous individual cells or fewer cells with numerous gene copies per cell. Gong and Marchetti (2019) found that gene copy numbers for 7 eukaryotic phytoplankton varied from 2 to 166. *Ostreococcus tauri* had 3.4 copies of the 18S gene across 13 strains, as opposed to the dinoflagellate *Symbiodinium kawagutii* at 160 copies, the latter likely reflecting the repetitive genomes of dinoflagellates in general (reviewed by Murray et al., 2016). A data compilation by Martin et al. (2022) reported literature values for gene copy number per cell for dinoflagellates and other flagellates (which encompassed the most groups that we report here), finding median values of 4919 and 5.23, respectively. Thus, our DNA results can be best be viewed as substantially overestimating the contribution of dinoflagellates relative to other groups.

In contrast, not all dinoflagellates have the pigment peridinin (∼2/3 of species studied by Zapata et al., 2012), which we used as a diagnostic proxy for them, and other work has shown that HPLC-peridinin based contributions are lower than parallel microscopy-based estimates (Latasa et al., 2022). Thus, the HPLC data likely underestimates their contribution to community biomass. Nevertheless, A-DINO and their indicator pigment PER did show good agreement in terms of their depth profiles (Fig. 6). The mean (± 1 standard deviation) EZ-integrated PER concentration for the Argo Basin was 0.5 ± 0.1 mg m^-2^, higher than that found for HOT (0.3 ± 0.1 mg m^-2^) or in the GoM (0.2 ± 0.1 mg m^-2^, Table 8), suggesting that PER-containing A-DINO are more prevalent in the Argo Basin than the other sites (Table 8).

Finally, microscopy-based abundance estimates for dinoflagellates are the most accurate, as this group is easily recognized visually. However, while the EPI-MICRO method distinguishes between cells with and without chlorophyll, the INVERT-MICRO method does not, whereas the IFCB method used here only enumerates chlorophyll-bearing cells. We found the IFCB 5-m samples had ∼10-times more dinoflagellate biomass than INVERT- and EPI-MICRO data (21 vs. 2-3.5 µg C L^-1^, respectively), although their average abundance was similar (6,023 ± 325 cells mL^-1^ for IFCB and 5,813 ± 901 cells mL^-1^ for INVERT-MICRO). This result might reflect the different depth horizons sampled (5-m vs. pooled 5-40 m), patchiness and small sample size from the IFCB method (∼5 mL) versus 350-450 mL for EPI- and INVERT-MICRO methods, or the different approaches used to estimate cell biovolumes (IFCB: Moberg and Sosik, 2012; EPI-MICRO: Yingling et al., 2025).

Additionally, most dinoflagellate taxa have the capacity for mixotrophy (Stoecker, 1999). Taken all together, these methods constrain the biomass of chlorophyll-bearing dinoflagellates, but mainly provide maximum and minimum estimates of their contributions to photosynthesis and mixotrophy in the system.

#### 4.4.2. Prymnesiophytes

Both HPLC and DNA methods showed PRYM as the dominant eukaryotic taxa in the EZ, as has been found elsewhere in oligotrophic waters (Shi et al., 2009; Liu et al., 2009; Cuvelier et al., 2010, Jardillier et al., 2010; Latasa et al., 2023). PRYM contribute to both the pico- and nano-sized eukaryotic groups (Liu et al., 2009, Lepère et al., 2009; Jardillier et al., 2010), and members of this group are also mixotrophic (Hartmann et al., 2012; Unrein et al., 2014; Anderson et al., 2018; Livanou et al., 2019). Indeed, the dominant PRYM genera found were *Chrysochromulina* and *Phaeocystis*, which both may be mixotrophic (Stoecker et al., 2006; Smith and Trimborn, 2024). PRYM and the indicator pigment HEX showed good agreement in their depth profiles (Fig. 4). PRYM had about 6-times more MV-CHL*a* in the lower EZ relative to the upper EZ (Table 4), agreeing with the clear increase in this group at the DCM from the DNA data (Fig. 9B). Mean (± 1 standard deviation) integrated HEX was 3.2 ± 0.8 mg m^-2^, compared to the same mean at HOT and much higher than in the GoM (1.5-1.8 mg m^-2^, Table 8).

#### 4.4.3. Pelagophytes

While not as high a percentage of *T*CHL*a* as HEX, BUT had one of the highest accessory pigment concentrations. As the indicator pigment for PELAG, it showed good agreement with this taxonomic group, increasing with depth (Fig. 4). Mean integrated BUT was similar in the Argo Basin to HOT (1.4 vs. 1.6 mg m^-2^, respectively), but about twice as high as in the GoM (0.6-0.7 mg m^-2^, Table 8). The DNA data showed that PELAG, specifically *P. calceolata*, were much more common at the DCM relative to the surface layer. The pigment data showed the same trend, with ∼9-times the amount of PELAG-associated MV-CHL*a* in the lower EZ, as well as a higher percentage of the MV-CHL*a* relative to the upper EZ (Table 4). PELAG were also found as an important contributor to the eukaryotic community in oligotrophic sites in the eastern North Pacific, with greater abundances in the DCM than at the surface, agreeing with our results (Dupont et al., 2015; Choi et al., 2020 Gúerin et al., 2022; Latasa et al., 2023). PELAG have also been found to be important mixotrophs, feeding on bacteria, in the ultra-oligotrophic eastern Mediterranean Sea (Livanou et al., 2019).

#### 4.4.4. Prasinophytes

The other dominant eukaryotic group was prasinophytes, which are part of the chlorophyte group, and all have MV-CHL*b*, but may differ in terms of other indicator pigments (Supp. Table S1). MV-CHL*b* was only 4-6% of *T*CHL*a* in the upper EZ, increasing to 11-16% of *T*CHL*a* in the lower EZ. PRAS-1 was mostly found in the upper EZ, whereas PRAS-3 (with PRAS pigment) was absent there but had a high concentration in the lower EZ. For DNA data that could be assigned to genus, the dominant DCM genera were *Ostreococcus*, with some *Bathycoccus*.

Unlike other groups, prasinophytes did not follow phytoclass assignments to their main indicator pigments well in the lower EZ; PRAS-1 appears to better track PRAS, whereas PRAS-3 appears to track MV-CHL*b*. This apparent mismatch between pigments and taxa with depth is likely an artifact of the pigment ratios optimization routine, matching a low concentration of PRAS pigment and a low PRAS pigment ratio (0.120) with a high MV-CHL*b* ratio (0.900) to PRAS-3, along with a lower MV-CHL*b* pigment ratio (0.214) for PRAS-1 (Supp. Table S1).

Our DNA data does not differentiate between prasinophyte groups, including PRAS-1 and PRAS-3, although the dominant genera found (*Ostreococcus* and *Bathycoccus*) contain PRAS. Like the pigment results, these data show an increase in prasinophytes at the DCM relative to the surface (Fig. 9B). However, the DNA data give more mean gene copies L^-1^ of prasinophytes in Cycles 1 and 3 vs Cycles 2 and 4, which is not reflected in the pigment data. Given the complexity of this group (containing many clades), it is likely that gene copies within these clades also vary, which might explain why those trends are not seen. Other research shows a generally good agreement between MV-CHLb and prasinophytes, so further examination of this issue is needed (Lampe et al., 2024).

#### 4.4.5. Cryptophytes and diatoms

Minor contributions from CRYPT and DIAT were also observed. CRYPT occurred mainly in the lower EZ, and are known to grow in lower light levels because they have good accessory pigments for light harvesting (Richardson, 2022). Other studies have found this distribution (Livanou et al., 2019). CRYPT is also a known mixotroph-containing group.

FUCO, the indicator pigment for DIAT, was low, and this pigment is also found in other, more abundant, groups (PELAG, PRYM: Supp. Table S1). This is likely why the FUCO profiles do not match the DIAT profiles well, especially near the DCM (Fig. 7). Integrated mean (± 1 standard deviation) FUCO was 0.5 ± 0.2 mg m^-2^, which was lower than HOT (0.7 ± 0.4 mg m^-2^) and higher than in the GoM (0.2-0.3 mg m^-2^, Table 8). According to EPI-MICRO data, diatoms were rarely present; more were detected using other methods (IFCB, pooled INVERT-MICRO and DNA), but none exceeded 6% of total autotrophic carbon. Most taxa were pennates (e.g., *Pseudo-nitzschia)*, although DNA data also indicated the presence of diverse centric taxa, including some detection of genera, *Hemiaulus* and *Rhi*zo*solenia*, that may harbor intracellular nitrogen-fixing symbionts.

### 4.5. Autotrophic carbon

Our analyses produced abundance and pigment concentrations, as opposed to carbon biomass. Using established literature values to convert abundance to carbon biomass for PRO and SYN, we derived estimates for these organisms’ contribution to EZ standing stock. Eukaryotic phytoplankton biomass was estimated from abundance (PICO-EUKs from FCM), and abundance plus biovolume from EPI-MICRO for NANO- and MICRO-EUKS. These combined data show that total EZ phytoplankton biomass ranged from 776 to 1604 mg C m^-2^, with the low end of the range essentially the same (775 mg C m^-2^) as found for the HOT site averaging over 277 cruises using completely different methods (i.e., total POC and ATP to estimate living carbon; Karl et al., 2022). Using methods very similar to those reported here, Pasulka et al. (2013) found HOT EZ integrated summer phytoplankton biomass of ∼1500 mg C m^-2^. This compares to the GoM, which had an average of 905 mg C m^-2^ (Selph et al. 2021). Thus, all these oligotrophic sites had relatively similar phytoplankton biomass. The elevated carbon content in Cycle 1 corresponds with the elevated satellite-derived NPP (NPP_Sat_, Kranz et al., this issue) and is potentially related to the storm mixing event before this cycle, either mixing deep CHL*a* into the upper EZ or stimulating CHLa growth with a nutrient infusion.

### 4.6. Carbon:chlorophyll a ratios

Due to its ease of measurement by both direct and indirect (satellite) means, CHL*a* is often used to estimate phytoplankton biomass. However, a cell’s CHL*a* is a small part of its carbon biomass, which is the desired currency for food web calculations and system productivity. To bridge the gap, C:CHL*a* ratios are used to determine carbon-based phytoplankton biomass. In this study, we used biovolumes from microscopy and an often-used biovolume:C conversion equation (Menden-Deuer and Lessard, 2000), along with literature estimates of prokaryotic carbon to derive phytoplankton carbon biomass.

We found that C:*T*CHL*a* was 120 overall, averaged for all populations and depths, with a higher value (173) in the upper EZ and a lower value (60) in the deeper EZ. These data follow expected trends - a higher C:*T*CHL*a* ratio under surface nitrogen limitation, high light and temperature, and decreasing C:*T*CHL*a* deeper in the water column under lower light and closer to the nitricine (Buck et al., 1996; Geider et al., 1997; Taylor et al., 1997; Graff et al., 2015; Marañón et al., 2003; Behrenfeld et al., 2016). In particular, the trends are the same as found in the open-ocean Gulf of Mexico (GoM), with an overall average of 117 ± 58 (Selph et al., 2021). However, while upper EZ values were also similar between the GoM and Argo Basin (171 and 173, respectively), lower EZ (DCM) values were much lower in the GoM (39 ± 16) vs. the Argo Basin (60 ± 6). This is likely because the nitracline (using the DCM depth as its proxy) was deeper in the GoM (107 ± 5 m) versus the Argo Basin (77 ± 2 m), which suggests higher carbon biomass relative to CHL*a* for the latter. At HOT, Pasulka et al. (2013) found C:CHL*a* ratios highest at the surface in summer (156 ± 157 or 108 ± 41, with or without sporadic contributions from *Trichodesmium*, respectively), with a much lower ratio in the deep EZ (34 ±6). These HOT estimates are lower than we found in the Argo Basin, although the deep values are similar to the GoM.

PRO C:DV-CHL*a* ratios were higher than the *T*CHL*a*-based ratios, which is also expected as PRO generally has a higher ratio than eukaryotic cells (Veldhuis and Kraay, 2004; Sathyendranath et al., 2009; Smyth et al., 2023). Despite differences in PRO carbon cell^-1^ conversion factors resulting in a large variance in literature estimates, the values found in this study are within those found in other oligotrophic habitats (Casey et al., 2013; Phongphattarawat et al., 2023).

## 5. Summary and conclusions

The Argo Basin phytoplankton community in austral summer 2022 was typical of oligotrophic, nitrogen-limited, warm, open-ocean regions. It was dominated by picoplankton, mainly PRO, PRYM, A-DINO, and prasinophytes in the upper EZ. Its EZ may be divided into 2 layers: an upper layer with those dominant members, and a lower layer with those groups plus PELAG and CRYPT. Prasinophytes switch from taxa without PRAS in the upper EZ, to those with PRAS in the lower EZ. DIAT was a minor component of the community. Nitrogen-fixing taxa were present, but at low abundance. Most of the dominant eukaryotic taxa in the Argo Basin have known mixotrophic members, and a higher percent of presumptive phagotrophs were found amongst the chlorophyll-bearing protists in the upper EZ than the lower EZ.

Similar abundances of PRO and SYN are found in the Argo Basis and HOT site, but overall *T*CHL*a* is lower, largely due to lower DV-CHL*a*, suggesting that DV-CHL*a* per PRO is lower in the Argo Basin. MV-CHL*a* is similar between HOT and the Argo Basin, as are concentrations of the main accessory pigments HEX and BUT, while PER is somewhat higher than at HOT. The GoM has lower PRO abundance but higher SYN abundance than the Argo Basin, but similar MV-CHL*a*:DV-CHL*a* ratios, leading to similar integrated *T*CHL*a*.

This is the first comprehensive description of this area, using multiple, complementary, independent methods for assessing phytoplankton taxonomic composition. The methods generally agreed, and where they did not, we use that information to constrain interpretations. For instance, A-DINO was an important group but the methods used to enumerate them all had their strengths and weaknesses, and together point to this group’s importance in oligotrophic regions.

## Acknowledgements

We thank the captain and crew of the R/V *Roger Revelle* for their excellent ship-based support, use of scientific collection equipment, and assistance with logistics. Research support was provided by the National Science Foundation Grants OCE-1756884 (to A.E.A.), OCE-1851381 (to K.E.S.), OCE-1851347 and OCE-1851558 (to M.R.L.), OCE-2332036 and OCE-2404504 (to S.A.K), National Ocean and Atmospheric Administration Grant NA19NOS4780181 (to A.E.A.), and Simons Foundation Collaboration on Principles of Microbial Ecosystems (PriME) Grant 970820 (to A.E.A). Seawater and plankton samples were collected under Australian Government permit AU-COM2021-520 and Australian Marine Parks permit PA2021-00062-2 issued by the Director of National Parks, Australia. The views expressed in this publication do not necessarily represent those of the Director of National Parks or the Australian Government.

## Author Statement

K.E.S. developed the protocols for measurements using flow cytometry. A.E.A. developed protocols for DNA extractions. K.E.S., R.H.L., N.Y., S.A.K. and M.R.L. conducted field sampling. K.E.S., R.H.L., N.Y., R.B. and S.A.K. analyzed results for flow cytometry, DNA, EPI-MICRO, INVERT-MICRO, and nutrients, respectively. R.H.L. and R. B. also analyzed results from IFCB. R.H.L. performed the DNA-based analyses. K.E.S. and R.H.L. wrote the manuscript, and all authors provided feedback on concepts in the manuscript, and provided edits of the manuscript.

## Supplementary Table and Figure Legends

**Supplementary Table S1.** Final pigment ratios for shallow (<50 m) and deep (≥50 m) samples, used in phytoclass program (v 2.0.0). a. Final shallow ratios; b. final deep ratios. Taxa are prymnesiophytes (PRYM), pelagophytes (PELAG), diatoms type 2 (DIAT), dinoflagellates (A-DINO), prasinophytes group 3 (PRAS-3, deep only), prasinophytes group 1 (PRAS-1), and cryptophytes (CRYPT). Pigments are partitioned as proportions of monovinyl chlorophyll *a* (MV-CHL*a*) using 19’-butanoly-fucoxanthin (BUT), fucoxanthin (FUCO), 19’-hexanoyl-fucoxanthin (HEX), peridinin (PER), prasinoxanthin (PRAS), neoxanthin (NEOX), violaxanthin (VIOL), lutein (LUT), monvinyl chlorophyll *b* (MV-CHL*b*), and allophycocyanin (ALLO).

**Supplementary Table S2.** Station locations (Latitude, °N; Longitude, °E), date/time (UTC) of sampling, sample gear, cycle, transect or Argo site, CTD or TM cast number, depth of sampling (m), and sample types collected. Sample gear: UW = underway ship system (2 m) or towed fish (1 m), NI = Niskin bottle and TM = Niskin-X bottle. Sample types: D = DNA, F = flow cytometry, H = HPLC pigments, M = epifluorescence microscopy, and C = CHL*a* (fluorometric).

**Supplementary Table S3.** Cycle 1 depth (m) profiles of pigments (ng L^-1^) from HPLC in CTD 11, 18, 25, 33, and 38. Allophycocyanin, violaxanthin, and lutien were all <4 ng L^-1^, so not shown; BDL = below detection limit.

**Supplementary Table S4.** Cycle 2 depth (m) profiles of pigments (ng L^-1^) from HPLC in CTD 48, 54, 61, 68, and 74. Allophycocyanin, violaxanthin, and lutien all had values <4 ng L^-1^, so not shown; BDL = below detection limit.

**Supplementary Table S5.** Cycle 3 depth (m) profiles of pigments (ng L^-1^) from HPLC in CTD 86, 93, 99, and 106. Allophycocyanin, violaxanthin, and lutien all had values <4 ng L^-1^, so not shown; BDL = below detection limit.

**Supplementary Table S6.** Cycle 4 depth (m) profiles of pigments (ng L^-1^) from HPLC in CTD 114, 121, and 128. Allophycocyanin, violaxanthin, and lutien all had values <4 ng L^-1^, so not shown; BDL = below detection limit.

**Supplementary Table S7.** Cycle 1 depth (m) profiles of taxa as contribution to monovinyl chlorophyll a (ng L^-1^) from HPLC in CTD 11, 18, 25, 33, and 38. *Synechococcus* (SYN) values are from flow cytometry (see text). All other taxa are from Phytoclass assignments and represent prymnesiophytes (PRYM), pelagophytes (PELAG), diatoms type 2 (DIAT), dinoflagellates (A-DINO), prasinophytes group 3 (PRAS-3), prasinophytes group 1 (PRAS-1) and cryptophytes (CRYPT).

**Supplementary Table S8.** Cycle 2 depth (m) profiles of taxa as contribution to monovinyl chlorophyll a (ng L^-1^) from HPLC in CTD 48, 54, 61, 68, and 74. *Synechococcus* (SYN) values are from flow cytometry (see text). All other taxa are from Phytoclass assignments and represent prymnesiophytes (PRYM), pelagophytes (PELAG), diatoms type 2 (DIAT), dinoflagellates (A-DINO), prasinophytes group 3 (PRAS-3), prasinophytes group 1 (PRAS-1) and cryptophytes (CRYPT).

**Supplementary Table S9.** Cycle 3 depth (m) profiles of taxa as contribution to monovinyl chlorophyll a (ng L^-1^) from HPLC in CTD 86, 93, 99, and 106. *Synechococcus* (SYN) values are from flow cytometry (see text). All other taxa are from Phytoclass assignments and represent prymnesiophytes (PRYM), pelagophytes (PELAG), diatoms type 2 (DIAT), dinoflagellates (A-DINO), prasinophytes group 3 (PRAS-3), prasinophytes group 1 (PRAS-1) and cryptophytes (CRYPT).

**Supplementary Table S10.** Cycle 4 depth (m) profiles of taxa as contribution to monovinyl chlorophyll a (ng L^-1^) from HPLC in CTD 114, 121, and 128. *Synechococcus* (SYN) values are from flow cytometry (see text). All other taxa are from Phytoclass assignments and represent prymnesiophytes (PRYM), pelagophytes (PELAG), diatoms type 2 (DIAT), dinoflagellates (A-DINO), prasinophytes group 3 (PRAS-3), prasinophytes group 1 (PRAS-1) and cryptophytes (CRYPT).

**Supplemental Figure S1.** Relationship between the sum of red fluorescence (SUM RF, abundance-weighted normalized chlorophyll fluorescence from flow cytometry) from all phytoplankton groups as a function of total CHLa (*T*CHL*a*, HPLC) in the same sample. Model II linear regression is: Sum RF = −289 + 6.93x*T*CHL*a*, r^2^ = 0.89. For details on how Sum RF is estimated, see Selph et al., 2021.

**Supplementary Figure S2**. Diversity of the eukaryotic phytoplankton ASVs (18S rRNA genes) between the mixed layer (blue) and DCM (green) among cycles, Argo stations, or transect stations. A) Alpha diversity expressed as the Shannon index. B) PCoA of Bray-Curtis dissimilarities.

**Supplementary Figure S3.** Genus-level DNA-based relative abundances among A-DINO within each cycle, Argo stations, or transect samples. Where genus-level assignment is not available, the lowest taxonomic level is shown (*Gymnodiniales* or *Dinophyceae*).

