## Supplementary material for "Phytoplankton community composition in the oligotrophic Argo Basin of the eastern Indian Ocean": Supp. Tables and Figures

**Supplementary Table S1.** Final pigment ratios for shallow (<50 m) and deep (≥50 m) samples, used in phytoclass program (v. 2.0.0). a. Final shallow ratios; b. final deep ratios. Taxa are prymnesiophytes (PRYM), pelagophytes (PELAG), diatoms type 2 (DIAT), dinoflagellates (A-DINO), prasinophytes group 3 (PRAS-3, deep only), prasinophytes group 1 (PRAS-1), and cryptophytes (CRYPT). Pigments are partitioned as proportions of monovinyl chlorophyll *a* (MV-CHL*a*) using 19'-butanoly-fucoxanthin (BUT), fucoxanthin (FUCO), 19'-hexanoyl-fucoxanthin (HEX), peridinin (PER), prasinoxanthin (PRAS), neoxanthin (NEOX), violaxanthin (VIOL), lutein (LUT), monvinyl chlorophyll *b* (MV-CHL*b*), and allophycocyanin (ALLO).

**a.** Final pigment ratios for shallow (<50 m) samples.

| TAXA | BUT | FUCO | HEX | PER | NEOX | VIOL | LUT | MV-CHL*b* | ALLO |
|---|---|---|---|---|---|---|---|---|---|
| **PRYM** | 0.213 | 0.016 | 1.107 | | | | | | |
| **PELAG** | 0.246 | 0.310 | | | | | | | |
| **DIAT** | | 0.605 | | | | | | | |
| **A-DINO** | | | | 0.348 | | | | | |
| **PRAS-1** | | | | | 0.012 | 0.064 | 0.107 | 0.381 | |
| **CRYPT** | | | | | | | | | 0.212 |

**b.** Final pigment ratios for deep (≥50 m) samples.

| TAXA | BUT | FUCO | HEX | PER | PRAS | NEOX | VIOL | LUT | MV-CHL*b* | ALLO |
|---|---|---|---|---|---|---|---|---|---|---|
| **PRYM** | 0.285 | 0.009 | 1.22 | | | | | | | |
| **PELAG** | 0.797 | 0.310 | | | | | | | | |
| **DIAT** | | 0.884 | | | | | | | | |
| **A-DINO** | | | | 0.403 | | | | | | |
| **PRAS-3** | | | | | 0.120 | 0.102 | 0.058 | 0.041 | 0.900 | |
| **PRAS-1** | | | | | | 0.050 | 0.141 | 0.107 | 0.214 | |
| **CRYPT** | | | | | | | | | | 0.219 |

**Supplementary Table S2.** Station locations (Latitude, °N; Longitude, °E), date/time (UTC) of sampling, sample gear, cycle, transect or Argo site, CTD or TM cast number, depth of sampling (m), and sample types collected. Sample gear: UW = underway ship system (2 m) or towed fish (1 m), NI = Niskin bottle and TM = Niskin-X bottle. Sample types: D = DNA, F = flow cytometry, H = HPLC pigments, M = epifluorescence microscopy, and C = CHL*a* (fluorometric).

| Latitude (°N) | Longitude (°E) | Date/Time (UTC) | Sample Gear | Cycle/ Transect/ ARGO | CTD/ TM No. | Depth(s) | Sample Type(s) |
|---|---|---|---|---|---|---|---|
| -10.2325 | 130.0771 | 1/26/22 23:00 | UW | T | - | 5 | D |
| -11.9174 | 130.0354 | 1/27/22 10:00 | UW | T | - | 5 | D |
| -12.4102 | 128.1925 | 1/29/22 0:00 | UW | T | - | 5 | D |
| -12.4008 | 126.2537 | 1/29/22 12:00 | UW | T | - | 5 | D |
| -13.0001 | 124.2044 | 1/30/22 1:00 | UW | T | - | 5 | D |
| -13.0403 | 122.3811 | 1/30/22 13:00 | UW | T | - | 5 | D |
| -13.1743 | 120.3293 | 1/31/22 1:00 | UW | T | - | 5 | D |
| -13.2744 | 119.4480 | 1/31/22 9:40 | NI | T | 2 | 11 | D |
| -13.2744 | 119.4480 | 1/31/22 9:40 | NI | T | 2 | 46 | D |
| -13.2536 | 119.1170 | 1/31/22 13:00 | UW | T | - | 5 | D |
| -13.2720 | 119.0985 | 1/31/22 14:20 | NI | T | 3 | 16 | D |
| -13.2720 | 119.0985 | 1/31/22 14:20 | NI | T | 3 | 55 | D |
| -13.3628 | 117.4655 | 2/1/22 1:00 | UW | T | - | 5 | D |
| -13.4800 | 115.3592 | 2/1/22 13:00 | UW | T | - | 5 | D |
| -13.5265 | 114.9475 | 2/1/22 17:00 | NI | T | 4 | 11 | D |
| -13.5265 | 114.9475 | 2/1/22 17:00 | NI | T | 4 | 76 | D |
| -13.5259 | 114.6410 | 2/1/22 21:00 | NI | T | 5 | 9 | D |
| -13.5259 | 114.6410 | 2/1/22 21:00 | NI | T | 5 | 79 | D |
| -13.5247 | 114.3142 | 2/2/22 0:20 | NI | T | 6 | 10 | D |
| -13.5247 | 114.3142 | 2/2/22 0:20 | NI | T | 6 | 85 | D |
| -13.5227 | 113.9555 | 2/2/22 3:35 | NI | T | 7 | 11 | D |
| -13.5227 | 113.9555 | 2/2/22 3:35 | NI | T | 7 | 80 | D |
| -15.3943 | 114.7054 | 2/3/22 2:00 | NI | T | 9 | 12 | D |
| -15.3943 | 114.7054 | 2/3/22 2:00 | NI | T | 9 | 76 | D |
| -15.4137 | 114.5983 | 2/3/22 12:00 | UW | T | - | 5 | D |
| -15.3494 | 114.5489 | 2/3/22 19:24 | NI | 1 | 11 | 5-90 | FHMC |
| -15.3505 | 114.5133 | 2/4/22 4:47 | NI | 1 | 14 | 10 | D |
| -15.3505 | 114.5133 | 2/4/22 4:47 | NI | 1 | 14 | 80 | D |
| -15.3747 | 114.4265 | 2/4/22 12:00 | UW | 1 | - | 5 | D |
| -15.3720 | 114.3663 | 2/4/22 17:56 | NI | 1 | 18 | 5-80 | FHMC |
| -15.3578 | 114.3374 | 2/5/22 4:15 | NI | 1 | 21 | 10 | D |

**Supplementary Table S2, continued**

| Latitude (°N) | Longitude (°E) | Date/Time (UTC) | Sample Gear | Cycle/ Transect/ ARGO | CTD/ TM No. | Depth(s) | Sample Type(s) |
|---|---|---|---|---|---|---|---|
| -15.3578 | 114.3374 | 2/5/22 4:15 | NI | 1 | 21 | 70 | D |
| -15.3944 | 114.2765 | 2/5/22 12:00 | UW | 1 | - | 5 | D |
| -15.3968 | 114.2331 | 2/5/22 18:02 | NI | 1 | 25 | 5-80 | FHMC |
| -15.4347 | 114.2033 | 2/6/22 5:40 | NI | 1 | 28 | 10 | D |
| -15.4347 | 114.2033 | 2/6/22 5:40 | NI | 1 | 28 | 80 | D |
| -15.4767 | 114.1806 | 2/6/22 12:00 | UW | 1 | - | 5 | D |
| -15.4683 | 114.1920 | 2/6/22 17:54 | NI | 1 | 33 | 5-65 | FHMC |
| -15.4995 | 114.1572 | 2/7/22 4:38 | NI | 1 | 36 | 10 | D |
| -15.4995 | 114.1572 | 2/7/22 4:38 | NI | 1 | 36 | 65 | D |
| -15.5597 | 114.1417 | 2/7/22 13:00 | UW | 1 | - | 5 | D |
| -15.5984 | 114.1351 | 2/7/22 19:06 | NI | 1 | 38 | 5-90 | FHC |
| -15.8138 | 114.5030 | 2/8/22 5:00 | NI | T | 39 | 10 | D |
| -15.8138 | 114.5030 | 2/8/22 5:00 | NI | T | 39 | 80 | D |
| -15.9850 | 114.7818 | 2/8/22 9:10 | NI | T | 40 | 10 | D |
| -15.9850 | 114.7818 | 2/8/22 9:10 | NI | T | 40 | 85 | D |
| -16.2421 | 115.0445 | 2/8/22 12:50 | NI | T | 41 | 10 | D |
| -16.2421 | 115.0445 | 2/8/22 12:50 | NI | T | 41 | 46 | D |
| -16.4706 | 115.2974 | 2/8/22 16:42 | NI | T | 42 | 17 | D |
| -16.9539 | 115.8064 | 2/9/22 1:00 | NI | T | 44 | 10 | D |
| -16.9539 | 115.8064 | 2/9/22 1:00 | NI | T | 44 | 65 | D |
| -17.1824 | 116.0338 | 2/9/22 4:30 | NI | T | 45 | 10 | D |
| -17.1824 | 116.0338 | 2/9/22 4:30 | NI | T | 45 | 70 | D |
| -16.7462 | 115.7089 | 2/9/22 19:02 | NI | 2 | 48 | 5-80 | FHMC |
| -16.7399 | 115.7618 | 2/10/22 5:30 | NI | 2 | 51 | 10 | D |
| -16.7399 | 115.7618 | 2/10/22 5:30 | NI | 2 | 51 | 80 | D |
| -16.8325 | 115.8468 | 2/10/22 12:00 | UW | 2 | - | 5 | D |
| -16.8171 | 115.8330 | 2/10/22 18:29 | NI | 2 | 54 | 5-80 | FHMC |
| -16.8328 | 115.9060 | 2/11/22 4:54 | NI | 2 | 57 | 10 | D |
| -16.8328 | 115.9060 | 2/11/22 4:54 | NI | 2 | 57 | 80 | D |
| -16.8442 | 115.9333 | 2/11/22 12:00 | UW | 2 | - | 5 | D |
| -16.8801 | 115.9441 | 2/11/22 18:01 | NI | 2 | 61 | 5-90 | FHMC |
| -16.9544 | 115.9932 | 2/12/22 5:20 | NI | 2 | 64 | 10 | D |
| -16.9544 | 115.9932 | 2/12/22 5:20 | NI | 2 | 64 | 80 | D |
| -16.9498 | 115.9842 | 2/12/22 5:00 | UW | 2 | - | 5 | D |
| -17.0368 | 116.0750 | 2/13/22 0:20 | UW | 2 | - | 5 | D |
| -16.9995 | 116.0730 | 2/12/22 18:01 | NI | 2 | 68 | 5-80 | FHMC |

**Supplementary Table S2, continued**

| Latitude (°N) | Longitude (°E) | Date/Time (UTC) | Sample Gear | Cycle/ Transect/ ARGO | CTD/ TM No. | Depth(s) | Sample Type(s) |
|---|---|---|---|---|---|---|---|
| -17.0610 | 116.1002 | 2/13/22 5:05 | NI | 2 | 71 | 10 | D |
| -17.0610 | 116.1002 | 2/13/22 5:05 | NI | 2 | 71 | 70 | D |
| -17.0691 | 116.1136 | 2/13/22 8:00 | UW | 2 | - | 5 | D |
| -17.1263 | 116.1493 | 2/13/22 19:00 | NI | 2 | 74 | 5-80 | FHC |
| -16.1701 | 115.6541 | 2/13/22 16:30 | UW | T | - | 5 | D |
| -16.0155 | 115.9087 | 2/15/22 7:55 | NI | T | 82 | 10 | D |
| -16.0155 | 115.9087 | 2/15/22 7:55 | NI | T | 82 | 75 | D |
| -15.9428 | 115.8287 | 2/15/22 18:26 | NI | 3 | 86 | 5-80 | FHMC |
| -15.9241 | 115.8308 | 2/16/22 8:25 | NI | 3 | 89 | 10 | D |
| -15.9241 | 115.8308 | 2/16/22 8:25 | NI | 3 | 89 | 70 | D |
| -15.9198 | 115.8294 | 2/16/22 5:50 | UW | 3 | - | 5 | D |
| -15.9156 | 115.8306 | 2/16/22 10:25 | UW | 3 | - | 5 | D |
| -15.9309 | 115.8586 | 2/16/22 16:00 | UW | 3 | - | 5 | D |
| -15.8494 | 115.8640 | 2/16/23 22:40 | UW | 3 | - | 5 | D |
| -15.8807 | 115.8381 | 2/16/22 18:05 | NI | 3 | 93 | 5-70 | FHMC |
| -15.8254 | 115.8291 | 2/17/22 4:30 | NI | 3 | 95 | 10 | D |
| -15.8254 | 115.8291 | 2/17/22 4:30 | NI | 3 | 95 | 60 | D |
| -15.8253 | 115.8291 | 2/17/22 6:20 | UW | 3 | - | 5 | D |
| -15.8066 | 115.8198 | 2/17/22 10:30 | UW | 3 | - | 5 | D |
| -15.7860 | 115.8295 | 2/17/22 16:45 | UW | 3 | - | 5 | D |
| -15.7641 | 115.7990 | 2/17/22 18:01 | NI | 3 | 99 | 5-60 | FHMC |
| -15.7033 | 115.7741 | 2/18/22 4:35 | NI | 3 | 102 | 10 | D |
| -15.7033 | 115.7741 | 2/18/22 4:35 | NI | 3 | 102 | 70 | D |
| -15.6003 | 115.7755 | 2/18/22 12:00 | UW | 3 | - | 5 | D |
| -15.6003 | 115.7757 | 2/18/22 19:08 | NI | 3 | 106 | 5-80 | FHC |
| -15.8781 | 116.5080 | 2/19/22 7:35 | NI | T | 107 | 10 | D |
| -15.8781 | 116.5080 | 2/19/22 7:35 | NI | T | 107 | 72 | D |
| -15.9129 | 117.0058 | 2/19/22 12:37 | NI | T | 108 | 10 | D |
| -15.9129 | 117.0058 | 2/19/22 12:37 | NI | T | 108 | 78 | D |
| -15.9955 | 117.9911 | 2/19/22 17:58 | NI | T | 110 | 70 | D |
| -15.8946 | 118.1368 | 2/20/22 11:30 | UW | T | - | 5 | D |
| -15.9011 | 118.1591 | 2/21/22 4:00 | UW | T | - | 5 | D |
| -15.8853 | 118.1423 | 2/20/22 18:18 | NI | 4 | 114 | 5-80 | FHMC |
| -15.8875 | 118.1320 | 2/21/22 4:20 | NI | 4 | 117 | 10 | D |
| -15.8875 | 118.1320 | 2/21/22 4:20 | NI | 4 | 117 | 80 | D |
| -15.9042 | 118.0938 | 2/21/22 19:50 | UW | 4 | - | 5 | D |

**Supplementary Table S2, continued**

| Latitude (°N) | Longitude (°E) | Date/Time (UTC) | Sample Gear | Cycle/ Transect/ ARGO | CTD/ TM No. | Depth(s) | Sample Type(s) |
|---|---|---|---|---|---|---|---|
| -15.9089 | 118.0893 | 2/21/22 18:07 | NI | 4 | 121 | 5-80 | FHMC |
| -15.9529 | 118.1058 | 2/22/22 4:30 | NI | 4 | 124 | 10 | D |
| -15.9529 | 118.1058 | 2/22/22 4:30 | NI | 4 | 124 | 80 | D |
| -15.9564 | 118.1251 | 2/22/22 12:10 | UW | 4 | - | 5 | D |
| -15.9392 | 118.1204 | 2/22/22 18:05 | NI | 4 | 128 | 5-80 | FHMC |
| -15.9483 | 118.0988 | 2/23/22 4:45 | NI | 4 | 131 | 10 | D |
| -15.9483 | 118.0988 | 2/23/22 4:45 | NI | 4 | 131 | 70 | D |
| -15.0000 | 119.3273 | 2/24/22 17:40 | UW | T | - | 5 | D |
| -15.0017 | 117.5019 | 2/25/22 8:15 | TM | ARGO | 4 | 10 | FDC |
| -15.0017 | 117.5019 | 2/25/22 8:15 | TM | ARGO | 4 | 76 | FDC |
| -15.0028 | 117.5028 | 2/25/22 8:30 | UW | T | - | 1 | D |
| -15.0014 | 115.4998 | 2/26/22 2:30 | UW | T | - | 5 | D |
| -15.0003 | 114.5032 | 2/26/22 11:30 | TM | ARGO | 7 | 10 | FDC |
| -15.0003 | 114.5032 | 2/26/22 11:30 | TM | ARGO | 7 | 65 | FDC |
| -15.0007 | 114.5011 | 2/26/22 11:30 | UW | T | - | 1 | D |
| -14.0011 | 114.4996 | 2/26/22 22:45 | UW | T | - | 1 | D |
| -13.0099 | 114.5022 | 2/27/22 11:00 | UW | T | - | 1 | D |
| -13.0099 | 114.5022 | 2/27/22 11:00 | TM | ARGO | 9 | 10 | FDC |
| -13.0099 | 114.5022 | 2/27/22 11:00 | TM | ARGO | 9 | 85 | FDC |
| -13.5086 | 116.5002 | 2/28/22 7:30 | TM | ARGO | 11 | 10 | FDC |
| -13.5086 | 116.5002 | 2/28/22 7:30 | TM | ARGO | 11 | 75 | FDC |
| -13.5086 | 116.5002 | 2/28/22 7:50 | UW | T | - | 1 | D |
| -13.5007 | 118.4931 | 3/1/22 0:15 | UW | T | - | 1 | D |
| -13.5053 | 119.4974 | 3/1/22 9:00 | TM | ARGO | 14 | 10 | FDC |
| -13.5053 | 119.4974 | 3/1/22 9:00 | TM | ARGO | 14 | 84 | FDC |
| -13.5054 | 119.4974 | 3/1/22 9:10 | UW | T | - | 1 | D |
| -13.6422 | 121.4878 | 3/2/22 10:30 | UW | T | - | 1 | D |
| -13.4964 | 121.4976 | 3/2/22 10:30 | TM | ARGO | 16 | 10 | FDC |
| -13.4964 | 121.4976 | 3/2/22 10:30 | TM | ARGO | 16 | 45 | FDC |
| -14.5508 | 121.2604 | 3/2/22 23:40 | UW | T | - | 1 | D |
| -15.1266 | 120.7123 | 3/3/22 4:00 | UW | T | - | 1 | D |
| -14.3996 | 122.0378 | 3/3/22 12:00 | UW | T | - | 1 | D |
| -14.2301 | 122.2467 | 3/3/25 13:15 | UW | T | - | 1 | D |
| -13.6218 | 122.6197 | 3/3/22 16:45 | UW | T | - | 1 | D |
| -12.4380 | 125.9776 | 3/4/22 11:00 | UW | T | - | 1 | D |
| -12.4126 | 128.7148 | 3/5/22 2:00 | UW | T | - | 1 | D |

**Supplementary Table S3.** Cycle 1 depth (m) profiles of pigments (ng $L^{-1}$) from HPLC in CTD 11, 18, 25, 33, and 38. Allophycocyanin, violaxanthin, and lutien were all <4 ng $L^{-1}$, so not shown; BDL = below detection limit.

| Depth (m) | MV-CHLa | DV-CHLa | MV-CHLb | ZEAX | HEX | BUT | PER | FUCO | PRAS | NEOX |
|---|---|---|---|---|---|---|---|---|---|---|
| **CTD 11** | | | | | | | | | | |
| 5 | 38.00 | 29.40 | 4.10 | 63.8 | 11.80 | 3.10 | 3.40 | 2.90 | BDL | 0.60 |
| 25 | 38.10 | 29.70 | 4.30 | 62.2 | 12.20 | 3.00 | 3.00 | 2.90 | BDL | 0.60 |
| 50 | 60.10 | 40.30 | 7.00 | 66.6 | 19.40 | 5.00 | 6.70 | 3.70 | BDL | 0.90 |
| 70 | 238.50 | 131.30 | 59.90 | 68.4 | 121.90 | 65.20 | 12.80 | 13.70 | 3.50 | 5.30 |
| 80 | 269.50 | 110.80 | 56.90 | 42.3 | 121.70 | 86.50 | 6.40 | 22.30 | 7.70 | 7.70 |
| 90 | 163.70 | 76.20 | 16.80 | 15.8 | 77.40 | 60.60 | 3.90 | 15.50 | 1.10 | 3.10 |
| **CTD 18*** | | | | | | | | | | |
| 20 | 40.60 | 33.00 | 3.80 | 62 | 11.50 | 3.10 | 4.20 | 2.70 | BDL | BDL |
| 40 | 53.10 | 40.40 | 5.40 | 61.3 | 17.80 | 4.30 | 5.10 | 3.30 | BDL | 0.60 |
| 60 | 159.10 | 99.70 | 19.30 | 56.5 | 78.50 | 31.60 | 7.60 | 8.50 | 1.30 | 2.00 |
| 70 | 282.80 | No data | 49.90 | 39.5 | 125.90 | 86.70 | 12.10 | 21.30 | 5.60 | 8.70 |
| 80 | 268.60 | 79.80 | 36.30 | 28.7 | 119.10 | 82.90 | 8.10 | 20.50 | 4.40 | 6.20 |
| **CTD 25*** | | | | | | | | | | |
| 5 | 45.10 | 57.50 | 4.50 | 63.8 | 15.00 | 3.40 | 3.90 | 2.90 | BDL | 0.60 |
| 20 | 48.70 | 61.80 | 4.50 | 66.9 | 17.40 | 4.20 | 4.20 | 3.30 | BDL | 0.70 |
| 40 | 52.40 | 63.90 | 5.20 | 68.2 | 18.70 | 4.30 | 4.80 | 3.40 | BDL | 0.70 |
| 50 | 76.00 | 105.50 | 6.80 | 74.7 | 31.70 | 7.30 | 6.60 | 4.80 | BDL | 0.90 |
| 60 | 118.30 | 119.00 | 12.80 | 68 | 58.70 | 15.00 | 10.20 | 5.70 | BDL | 1.50 |
| **CTD 33** | | | | | | | | | | |
| 5 | 45.70 | 55.30 | 3.60 | 75.4 | 14.00 | 3.30 | 6.00 | 2.90 | BDL | BDL |
| 20 | 44.50 | 58.30 | 3.80 | 74.6 | 14.30 | 3.40 | 3.90 | 2.80 | BDL | BDL |
| 30 | 58.00 | 70.40 | 5.00 | 68.3 | 20.30 | 4.80 | 4.70 | 3.50 | BDL | 0.60 |
| 40 | 84.30 | 116.00 | 6.20 | 62.9 | 33.30 | 7.50 | 5.90 | 4.50 | BDL | 0.90 |
| 50 | 151.90 | 145.40 | 14.00 | 63.3 | 71.20 | 22.70 | 10.00 | 6.50 | BDL | 1.30 |
| 65 | 257.60 | 113.60 | 47.10 | 44.6 | 131.90 | 71.50 | 12.40 | 16.20 | 3.20 | 6.30 |
| **CTD 38** | | | | | | | | | | |
| 5 | 31.00 | 40.90 | 3.00 | 68.8 | 7.70 | 2.00 | 2.70 | 2.00 | BDL | BDL |
| 20 | 35.00 | 42.20 | 3.90 | 70.5 | 9.10 | 2.30 | 3.60 | 2.30 | BDL | BDL |
| 40 | 47.40 | 51.40 | 5.80 | 76.8 | 14.10 | 4.00 | 4.50 | 2.90 | BDL | 0.70 |
| 60 | 69.80 | 85.50 | 9.30 | 87.4 | 27.00 | 8.80 | 5.50 | 4.10 | BDL | 0.70 |
| 80 | 231.00 | 109.30 | 51.40 | 56.5 | 114.00 | 51.90 | 9.60 | 11.60 | 3.50 | 4.50 |
| 90 | 303.40 | 93.80 | 73.70 | 41.5 | 133.20 | 84.00 | 9.10 | 19.00 | 6.60 | 9.40 |

. **CTD 18 has no 5 m sample; CTD 25 has no 80 m sample*

**Supplementary Table S4.** Cycle 2 depth (m) profiles of pigments (ng $L^{-1}$) from HPLC in CTD 48, 54, 61, 68, and 74. Allophycocyanin, violaxanthin, and lutien all had values <4 ng $L^{-1}$, so not shown; BDL = below detection limit.

| Depth (m) | MV-CHLa | DV-CHLa | MV-CHLb | ZEAX | HEX | BUT | PER | FUCO | PRAS | NEOX |
|---|---|---|---|---|---|---|---|---|---|---|
| **CTD 48** | | | | | | | | | | |
| 5 | 79.50 | 46.40 | 9.80 | 77.8 | 20.30 | 6.80 | 8.30 | 4.50 | BLD | 1.00 |
| 20 | 44.50 | 32.90 | 5.80 | 79.3 | 11.50 | 3.70 | 3.50 | 2.90 | BLD | 0.70 |
| 40 | 37.80 | 40.40 | 3.40 | 65.8 | 8.00 | 2.60 | 2.70 | 2.40 | BLD | 0.40 |
| 60 | 133.30 | 114.30 | 17.80 | 79.6 | 47.30 | 14.80 | 12.40 | 6.50 | BLD | 2.00 |
| 70 | 254.20 | 178.60 | 46.30 | 84.0 | 89.60 | 51.10 | 12.40 | 18.20 | 3.90 | 4.20 |
| 80 | 310.60 | 141.70 | 43.00 | 52.0 | 121.20 | 89.70 | 12.80 | 21.50 | 4.20 | 6.00 |
| **CTD 54** | | | | | | | | | | |
| 5 | 34.70 | 32.40 | 3.00 | 60.8 | 7.50 | 2.50 | 2.30 | 2.30 | BLD | BLD |
| 20 | 35.80 | 26.70 | 4.60 | 70.9 | 8.00 | 2.40 | 3.00 | 1.90 | BLD | BLD |
| 40 | 62.70 | 43.90 | 7.30 | 69.7 | 15.80 | 5.10 | 6.20 | 3.60 | BLD | 0.70 |
| 60 | 95.80 | 72.20 | 11.20 | 69.8 | 30.30 | 10.00 | 6.40 | 5.10 | 0.50 | 1.10 |
| 70 | 145.60 | 103.60 | 17.10 | 79.5 | 52.40 | 16.60 | 14.80 | 7.20 | 1.40 | 2.40 |
| 80 | 247.20 | 195.40 | 52.80 | 103.7 | 78.10 | 54.90 | 7.80 | 14.00 | 7.00 | 6.10 |
| **CTD 61** | | | | | | | | | | |
| 5 | 34.90 | 34.50 | 2.70 | 66.4 | 7.40 | 2.50 | 2.30 | 3.30 | BLD | BLD |
| 20 | 37.40 | 35.00 | 4.00 | 83.3 | 9.50 | 2.70 | 2.90 | 2.20 | BLD | 0.60 |
| 40 | 43.20 | 31.40 | 4.90 | 81 | 11.90 | 3.20 | 4.20 | 2.60 | BLD | 0.70 |
| 60 | 80.60 | 60.00 | 8.30 | 77 | 24.40 | 7.60 | 8.60 | 3.90 | BLD | 0.90 |
| 80 | 170.40 | 121.40 | 30.70 | 86.4 | 61.70 | 26.30 | 10.40 | 8.10 | 2.60 | 3.10 |
| 90 | 371.30 | 177.40 | 63.80 | 67.2 | 151.70 | 106.30 | 12.80 | 26.70 | 6.80 | 9.40 |
| **CTD 68*** | | | | | | | | | | |
| 20 | 38.30 | 31.80 | 4.60 | 90.1 | 10.80 | 2.80 | 3.50 | 1.90 | BLD | 0.60 |
| 40 | 46.70 | 34.00 | 4.50 | 84.6 | 13.10 | 3.60 | 4.70 | 2.50 | BLD | 0.60 |
| 60 | 128.40 | 89.70 | 15.50 | 80.8 | 52.80 | 14.00 | 9.00 | 6.50 | 0.80 | 1.50 |
| 70 | 322.80 | 143.60 | 73.20 | 83.2 | 143.60 | 79.10 | 15.60 | 19.50 | 7.60 | 7.60 |
| 80 | 328.40 | 143.10 | 78.90 | 70.6 | 123.10 | 100.70 | 11.30 | 21.60 | 6.70 | 10.30 |
| **CTD 74** | | | | | | | | | | |
| 5 | 32.80 | 36.40 | 2.90 | 65.3 | 8.20 | 2.30 | 2.40 | 2.30 | BLD | BLD |
| 20 | 36.90 | 29.60 | 3.40 | 89.6 | 9.10 | 2.30 | 3.40 | 1.70 | BLD | BLD |
| 40 | 79.00 | 50.10 | 7.60 | 84.4 | 26.00 | 6.80 | 8.00 | 4.40 | BLD | 0.90 |
| 60 | 279.10 | 121.40 | 48.30 | 77.5 | 136.20 | 56.20 | 15.40 | 14.20 | 5.40 | 5.10 |
| 70 | 313.50 | 126.90 | 56.90 | 59 | 127.80 | 97.70 | 9.30 | 20.40 | 5.60 | 7.60 |
| 80 | 261.70 | 107.80 | 25.20 | 38 | 117.00 | 84.70 | 9.20 | 18.80 | 2.90 | 4.70 |

. **CTD 68 has no 5 m sample*

**Supplementary Table S5.** Cycle 3 depth (m) profiles of pigments (ng $L^{-1}$) from HPLC in CTD 86, 93, 99, and 106. Allophycocyanin, violaxanthin, and lutien all had values <4 ng $L^{-1}$, so not shown; BDL = below detection limit.

| Depth (m) | MV-CHLa | DV-CHLa | MV-CHLb | ZEAX | HEX | BUT | PER | FUCO | PRAS | NEOX |
|---|---|---|---|---|---|---|---|---|---|---|
| **CTD 86** | | | | | | | | | | |
| 5 | 39.50 | 38.80 | 2.60 | 76.5 | 9.10 | 2.70 | 2.90 | 2.20 | BLD | BLD |
| 20 | 49.80 | 35.30 | 4.00 | 78.5 | 13.40 | 3.60 | 5.50 | 2.50 | BLD | BLD |
| 40 | 71.20 | 45.40 | 7.40 | 83.3 | 22.40 | 6.50 | 8.80 | 3.90 | BLD | 0.90 |
| 60 | 215.80 | 134.60 | 31.60 | 93.4 | 88.90 | 35.10 | 11.60 | 16.50 | 3.00 | 4.00 |
| 70 | 297.40 | 172.20 | 45.20 | 95.5 | 132.50 | 57.60 | 15.10 | 24.60 | 4.20 | 6.00 |
| 80 | 213.00 | 158.00 | 42.60 | 102.8 | 97.20 | 31.90 | 12.90 | 8.80 | 2.10 | 4.40 |
| **CTD 93** | | | | | | | | | | |
| 5 | 40.20 | 39.50 | 2.70 | 77.5 | 9.20 | 2.60 | 2.80 | 2.10 | BLD | BLD |
| 20 | 50.90 | 36.50 | 4.00 | 79.4 | 13.50 | 3.70 | 5.50 | 2.50 | BLD | BLD |
| 40 | 71.80 | 45.90 | 7.30 | 84.5 | 22.60 | 6.50 | 8.80 | 3.90 | BLD | 0.90 |
| 50 | 218.10 | 136.90 | 33.40 | 93.5 | 90.60 | 37.20 | 10.60 | 18.00 | 3.70 | 5.00 |
| 60 | 284.60 | 194.40 | 49.20 | 96.2 | 133.60 | 57.80 | 15.10 | 24.60 | 4.10 | 5.80 |
| 70 | 212.10 | 163.80 | 47.00 | 102.7 | 97.30 | 32.10 | 13.10 | 8.90 | 2.40 | 4.60 |
| **CTD 99** | | | | | | | | | | |
| 5 | 34.50 | 34.80 | 3.50 | 69 | 9.20 | 2.30 | 2.70 | 2.00 | BLD | BLD |
| 20 | 47.60 | 31.00 | 3.70 | 81.7 | 13.00 | 3.30 | 4.80 | 2.40 | BLD | 0.60 |
| 30 | 59.50 | 36.80 | 5.10 | 79.7 | 19.20 | 4.70 | 7.30 | 3.10 | BLD | 0.70 |
| 40 | 76.40 | 49.50 | 8.10 | 85.1 | 28.60 | 6.80 | 9.50 | 3.70 | BLD | 1.00 |
| 50 | 118.10 | 94.90 | 14.00 | 88.4 | 48.00 | 13.40 | 9.70 | 6.30 | BLD | 1.70 |
| 60 | 276.10 | 93.70 | 49.80 | 51.8 | 133.40 | 88.60 | 11.10 | 19.00 | 3.50 | BLD |
| **CTD 106** | | | | | | | | | | |
| 5 | 36.50 | 35.20 | 3.50 | 60.1 | 10.40 | 2.70 | 3.50 | 2.20 | BLD | 0.60 |
| 20 | 46.80 | 29.30 | 4.40 | 73.3 | 11.60 | 2.70 | 5.90 | 1.90 | BLD | 0.70 |
| 40 | 71.20 | 46.80 | 6.40 | 71.1 | 24.50 | 6.80 | 8.80 | 4.30 | BLD | 1.10 |
| 60 | 130.10 | 69.40 | 17.90 | 66.4 | 65.10 | 18.30 | 14.40 | 6.80 | BLD | 1.90 |
| 70 | 277.80 | 109.30 | 56.20 | 57.4 | 141.60 | 71.00 | 17.40 | 19.60 | 3.50 | 7.60 |
| 80 | 308.70 | 123.50 | 58.20 | 48.2 | 124.00 | 89.50 | 10.00 | 26.50 | 8.10 | BLD |

**Supplementary Table S6.** Cycle 4 depth (m) profiles of pigments (ng $L^{-1}$) from HPLC in CTD 114, 121, and 128. Allophycocyanin, violaxanthin, and lutien all had values <4 ng $L^{-1}$, so not shown; BDL = below detection limit.

| Depth (m) | MV-CHLa | DV-CHLa | MV-CHLb | ZEAX | HEX | BUT | PER | FUCO | PRAS | NEOX |
|---|---|---|---|---|---|---|---|---|---|---|
| **CTD 114** | | | | | | | | | | |
| 5 | 32.20 | 34.70 | 2.70 | 58.2 | 10.80 | 2.10 | 3.20 | 1.60 | BLD | BLD |
| 20 | 36.10 | 30.60 | 3.60 | 67.7 | 11.40 | 2.50 | 3.90 | 1.90 | BLD | 0.70 |
| 40 | 48.60 | 30.10 | 4.20 | 73.1 | 18.20 | 3.60 | 6.00 | 2.10 | BLD | BLD |
| 55 | 70.20 | 51.00 | 7.40 | 80.6 | 29.20 | 6.30 | 7.20 | 3.90 | BLD | 0.80 |
| 70 | 131.50 | 87.60 | 47.90 | 81.8 | 71.80 | 18.20 | 10.10 | 6.00 | 0.90 | BLD |
| 80 | 263.70 | 127.90 | 86.50 | 71.5 | 124.80 | 73.30 | 14.70 | 15.50 | 3.40 | BLD |
| **CTD 121** | | | | | | | | | | |
| 5 | 33.50 | 28.50 | 2.90 | 57 | 11.30 | 2.40 | 2.90 | 1.90 | BLD | BLD |
| 20 | 36.00 | 24.20 | 3.30 | 61.9 | 11.50 | 2.40 | 4.10 | 1.60 | BLD | 0.50 |
| 30 | 41.90 | 25.60 | 3.10 | 60.6 | 14.90 | 3.30 | 4.30 | 2.30 | BLD | BLD |
| 40 | 49.10 | 33.60 | 4.50 | 65.1 | 20.70 | 4.10 | 5.20 | 2.60 | BLD | 0.70 |
| 55 | 87.20 | 65.30 | 8.40 | 75.1 | 40.50 | 8.30 | 8.60 | 4.10 | BLD | 1.10 |
| 80 | 256.20 | 105.90 | 57.30 | 49.8 | 126.20 | 77.70 | 10.70 | 17.30 | 4.80 | 10.50 |
| **CTD 128** | | | | | | | | | | |
| 5 | 30.20 | 31.60 | 2.50 | 54.5 | 10.60 | 2.00 | 3.40 | 1.80 | BLD | BLD |
| 20 | 34.30 | 25.70 | 3.30 | 59 | 11.30 | 2.30 | 4.00 | 1.70 | BLD | BLD |
| 30 | 42.90 | 24.10 | 3.50 | 59.2 | 14.10 | 3.50 | 5.30 | 2.50 | BLD | 0.60 |
| 50 | 71.00 | 51.70 | 7.40 | 68.4 | 28.90 | 6.70 | 8.60 | 3.50 | BLD | 1.00 |
| 70 | 133.80 | 86.50 | 33.50 | 62.9 | 76.30 | 19.50 | 10.60 | 5.80 | 0.50 | BLD |
| 80 | 253.10 | 107.90 | 68.20 | 48.8 | 120.70 | 68.00 | 15.50 | 13.70 | 3.00 | BLD |

**Supplementary Table S7.** Cycle 1 depth (m) profiles of taxa as contribution to monovinyl chlorophyll a (ng $L^{-1}$) from HPLC in CTD 11, 18, 25, 33, and 38. *Synechococcus* (SYN) values are from flow cytometry (see text). All other taxa are from Phytoclass assignments and represent prymnesiophytes (PRYM), pelagophytes (PELAG), diatoms type 2 (DIAT), dinoflagellates (A-DINO), prasinophytes group 3 (PRAS-3), prasinophytes group 1 (PRAS-1) and cryptophytes (CRYPT).

| Depth (m) | SYN | PRYM | PELAG | DIAT | A-DINO | PRAS-3 | PRAS-1 | CRYPT |
|---|---|---|---|---|---|---|---|---|
| **CTD 11** | | | | | | | | |
| 5 | 3.88 | 10.21 | 3.21 | 2.68 | 9.34 | 0.00 | 8.68 | 0.00 |
| 25 | 4.06 | 10.76 | 2.57 | 3.08 | 8.41 | 0.00 | 9.22 | 0.00 |
| 50 | 5.66 | 18.52 | 0.67 | 4.39 | 19.15 | 7.00 | 4.52 | 0.19 |
| 70 | 22.44 | 88.41 | 37.49 | 0.00 | 27.62 | 53.17 | 0.00 | 9.37 |
| 80 | 1.66 | 108.20 | 74.88 | 0.00 | 17.02 | 67.34 | 0.00 | 0.40 |
| 90 | 0.31 | 69.41 | 52.80 | 0.00 | 10.30 | 19.16 | 1.38 | 10.35 |
| **CTD 18*** | | | | | | | | |
| 20 | 4.17 | 10.39 | 3.59 | 2.35 | 12.07 | 0.00 | 8.04 | 0.00 |
| 40 | 4.78 | 15.91 | 3.49 | 3.19 | 14.51 | 0.00 | 11.21 | 0.00 |
| 60 | 19.74 | 70.33 | 18.03 | 3.42 | 20.56 | 21.60 | 1.17 | 4.24 |
| 70 | ** | 110.77 | 70.95 | 0.00 | 31.49 | 58.79 | 0.00 | 10.80 |
| 80 | 1.14 | 113.22 | 73.48 | 0.00 | 22.84 | 46.37 | 0.00 | 11.55 |
| **CTD 25*** | | | | | | | | |
| 5 | 6.22 | 13.20 | 2.01 | 3.30 | 10.91 | 0.00 | 9.47 | 0.00 |
| 20 | 6.83 | 14.85 | 3.24 | 3.11 | 11.37 | 0.00 | 9.29 | 0.00 |
| 40 | 7.10 | 15.79 | 2.63 | 3.50 | 12.85 | 0.00 | 10.54 | 0.00 |
| 50 | 14.78 | 28.07 | 0.00 | 5.56 | 17.67 | 6.58 | 3.22 | 0.11 |
| 60 | 16.80 | 49.75 | 1.63 | 5.56 | 26.10 | 12.38 | 2.71 | 3.36 |
| **CTD 33** | | | | | | | | |
| 5 | 4.48 | 12.04 | 2.32 | 3.06 | 16.41 | 0.00 | 7.38 | 0.00 |
| 20 | 4.58 | 13.63 | 2.76 | 3.11 | 11.87 | 0.00 | 8.50 | 0.05 |
| 30 | 5.65 | 19.35 | 3.80 | 3.64 | 14.31 | 0.00 | 11.17 | 0.07 |
| 40 | 12.66 | 31.23 | 4.55 | 4.57 | 17.67 | 0.00 | 13.56 | 0.06 |
| 50 | 26.59 | 66.71 | 8.60 | 4.66 | 28.23 | 15.06 | 1.67 | 0.39 |
| 65 | 12.04 | 107.78 | 49.26 | 0.00 | 30.43 | 49.12 | 0.00 | 8.98 |
| **CTD 38** | | | | | | | | |
| 5 | 2.55 | 7.80 | 2.35 | 2.29 | 8.75 | 0.00 | 7.19 | 0.08 |
| 20 | 2.90 | 8.38 | 2.26 | 2.50 | 10.56 | 0.00 | 8.39 | 0.01 |
| 40 | 3.33 | 12.21 | 5.00 | 1.72 | 12.37 | 0.00 | 12.78 | 0.00 |
| 60 | 12.51 | 24.62 | 3.44 | 3.67 | 15.09 | 9.68 | 0.66 | 0.14 |
| 80 | 16.23 | 94.53 | 31.93 | 1.10 | 24.05 | 52.51 | 0.00 | 10.64 |
| 90 | 4.33 | 117.30 | 63.44 | 0.00 | 23.60 | 81.62 | 0.00 | 13.12 |

**CTD 18 has no 5 m sample; CTD 25 has no 80 m sample*

***No DV-CHLa, so no way to calculate SYN contribution to MV-CHLa*

**Supplementary Table S8.** Cycle 2 depth (m) profiles of taxa as contribution to monovinyl chlorophyll a (ng $L^{-1}$) from HPLC in CTD 48, 54, 61, 68, and 74. *Synechococcus* (SYN) values are from flow cytometry (see text). All other taxa are from Phytoclass assignments and represent prymnesiophytes (PRYM), pelagophytes (PELAG), diatoms type 2 (DIAT), dinoflagellates (A-DINO), prasinophytes group 3 (PRAS-3), prasinophytes group 1 (PRAS-1) and cryptophytes (CRYPT).

| Depth (m) | SYN | PRYM | PELAG | DIAT | A-DINO | PRAS-3 | PRAS-1 | CRYPT |
|---|---|---|---|---|---|---|---|---|
| **CTD 48** | | | | | | | | |
| 5 | 5.63 | 17.64 | 11.31 | 0.90 | 22.93 | 0.00 | 21.10 | 0.00 |
| 20 | 3.72 | 10.55 | 6.12 | 1.45 | 10.23 | 0.00 | 12.42 | 0.01 |
| 40 | 6.95 | 7.59 | 4.51 | 1.65 | 8.18 | 0.00 | 8.89 | 0.04 |
| 60 | 37.49 | 35.06 | 4.24 | 4.83 | 27.91 | 14.87 | 4.12 | 4.78 |
| 70 | 73.38 | 64.36 | 33.34 | 5.81 | 27.01 | 42.32 | 0.00 | 7.97 |
| 80 | 2.92 | 120.68 | 81.03 | 0.00 | 37.03 | 54.12 | 0.00 | 14.82 |
| **CTD 54** | | | | | | | | |
| 5 | 6.12 | 7.53 | 4.75 | 1.58 | 7.40 | 0.00 | 7.25 | 0.07 |
| 20 | 3.36 | 7.81 | 3.76 | 1.25 | 9.35 | 0.00 | 10.21 | 0.06 |
| 40 | 8.06 | 13.73 | 8.04 | 1.24 | 17.12 | 0.00 | 14.51 | 0.00 |
| 60 | 22.58 | 27.62 | 4.03 | 4.67 | 17.54 | 11.49 | 4.69 | 3.19 |
| 70 | 34.65 | 39.98 | 5.07 | 5.41 | 34.24 | 15.63 | 6.95 | 3.67 |
| 80 | 62.98 | 61.11 | 41.70 | 0.00 | 18.18 | 54.86 | 0.00 | 8.37 |
| **CTD 61** | | | | | | | | |
| 5 | 3.38 | 7.87 | 5.10 | 3.55 | 7.87 | 0.00 | 7.01 | 0.13 |
| 20 | 4.03 | 9.30 | 3.82 | 1.73 | 9.08 | 0.00 | 9.38 | 0.06 |
| 40 | 4.35 | 10.73 | 3.67 | 2.12 | 12.05 | 0.00 | 10.27 | 0.00 |
| 60 | 12.83 | 23.01 | 2.70 | 3.84 | 24.30 | 8.73 | 2.39 | 2.80 |
| 80 | 50.99 | 45.46 | 13.43 | 3.10 | 23.24 | 28.90 | 1.43 | 3.86 |
| 90 | 2.68 | 143.75 | 95.28 | 0.00 | 36.01 | 78.68 | 0.00 | 14.90 |
| **CTD 68** | | | | | | | | |
| 20 | 4.05 | 9.88 | 2.95 | 1.40 | 10.19 | 0.00 | 9.81 | 0.01 |
| 40 | 4.38 | 12.26 | 4.52 | 1.64 | 14.02 | 0.00 | 9.85 | 0.03 |
| 60 | 31.08 | 43.80 | 2.08 | 6.25 | 22.55 | 15.10 | 3.84 | 3.71 |
| 70 | 38.81 | 110.07 | 53.49 | 0.80 | 36.17 | 73.17 | 0.00 | 10.29 |
| 80 | 4.07 | 116.53 | 79.06 | 0.00 | 29.81 | 89.32 | 0.00 | 9.62 |
| **CTD 74** | | | | | | | | |
| 5 | 4.47 | 8.15 | 3.21 | 2.31 | 7.64 | 0.00 | 6.96 | 0.07 |
| 20 | 3.32 | 9.52 | 2.57 | 1.68 | 11.38 | 0.00 | 8.32 | 0.12 |
| 40 | 7.50 | 23.17 | 7.16 | 2.89 | 22.68 | 0.00 | 15.60 | 0.00 |
| 60 | 40.56 | 107.56 | 29.40 | 4.07 | 36.77 | 50.37 | 0.00 | 10.36 |
| 70 | 8.00 | 125.27 | 77.43 | 0.00 | 25.62 | 67.93 | 0.00 | 9.25 |
| 80 | 3.01 | 116.58 | 71.61 | 0.00 | 26.34 | 32.89 | 0.00 | 11.28 |

**CTD 68 has no 5 m sample*

**Supplementary Table S9.** Cycle 3 depth (m) profiles of taxa as contribution to monovinyl chlorophyll a (ng $L^{-1}$) from HPLC in CTD 86, 93, 99, and 106. *Synechococcus* (SYN) values are from flow cytometry (see text). All other taxa are from Phytoclass assignments and represent prymnesiophytes (PRYM), pelagophytes (PELAG), diatoms type 2 (DIAT), dinoflagellates (A-DINO), prasinophytes group 3 (PRAS-3), prasinophytes group 1 (PRAS-1) and cryptophytes (CRYPT).

| Depth (m) | SYN | PRYM | PELAG | DIAT | A-DINO | PRAS-3 | PRAS-1 | CRYPT |
|---|---|---|---|---|---|---|---|---|
| **CTD 86** | | | | | | | | |
| 5 | 5.22 | 9.12 | 4.25 | 1.60 | 9.31 | 0.00 | 7.83 | 2.16 |
| 20 | 5.03 | 12.79 | 4.36 | 1.79 | 16.74 | 0.00 | 9.03 | 0.06 |
| 40 | 6.29 | 18.53 | 8.13 | 1.25 | 23.11 | 0.00 | 13.90 | 0.00 |
| 60 | 46.46 | 69.09 | 17.02 | 11.05 | 27.27 | 30.51 | 8.54 | 5.87 |
| 70 | 63.43 | 100.05 | 30.80 | 13.88 | 34.50 | 44.29 | 4.11 | 6.34 |
| 80 | 46.70 | 72.83 | 10.51 | 4.68 | 29.26 | 38.92 | 4.12 | 5.96 |
| **CTD 93** | | | | | | | | |
| 5 | 3.97 | 10.40 | 4.19 | 1.90 | 10.22 | 0.00 | 9.31 | 0.21 |
| 20 | 4.37 | 13.33 | 4.87 | 1.66 | 17.33 | 0.00 | 9.24 | 0.10 |
| 40 | 6.14 | 18.90 | 8.07 | 1.34 | 23.37 | 0.00 | 13.98 | 0.00 |
| 50 | 26.09 | 77.70 | 20.91 | 13.10 | 27.45 | 36.41 | 10.55 | 5.89 |
| 60 | 62.96 | 94.58 | 28.90 | 13.07 | 32.37 | 45.08 | 1.57 | 6.08 |
| 70 | 6.56 | 86.43 | 12.61 | 5.55 | 35.10 | 49.93 | 7.70 | 8.22 |
| **CTD 99** | | | | | | | | |
| 5 | 5.92 | 7.95 | 2.05 | 1.91 | 7.41 | 0.00 | 7.03 | 2.23 |
| 20 | 3.94 | 13.13 | 3.61 | 2.23 | 15.50 | 0.00 | 9.07 | 0.12 |
| 30 | 5.31 | 16.83 | 3.93 | 2.52 | 20.35 | 0.00 | 10.56 | 0.00 |
| 40 | 6.89 | 23.07 | 4.65 | 2.47 | 24.28 | 0.00 | 15.04 | 0.00 |
| 50 | 25.80 | 38.37 | 2.64 | 5.63 | 23.45 | 11.90 | 6.79 | 3.51 |
| 60 | 14.51 | 113.40 | 62.19 | 0.00 | 27.25 | 47.71 | 1.15 | 9.89 |
| **CTD 106** | | | | | | | | |
| 5 | 6.53 | 8.53 | 2.57 | 1.77 | 9.10 | 0.00 | 7.99 | 0.00 |
| 20 | 3.89 | 10.98 | 1.98 | 1.99 | 17.79 | 0.00 | 10.13 | 0.05 |
| 40 | 6.39 | 20.09 | 7.67 | 2.00 | 22.89 | 0.00 | 12.16 | 0.00 |
| 60 | 29.71 | 43.45 | 3.13 | 4.76 | 29.25 | 13.50 | 3.35 | 2.97 |
| 70 | 27.39 | 103.06 | 42.37 | 3.87 | 38.35 | 52.79 | 0.00 | 9.98 |
| 80 | 3.15 | 111.80 | 82.77 | 2.60 | 27.25 | 65.06 | 0.00 | 16.07 |

**Supplementary Table S10.** Cycle 4 depth (m) profiles of taxa as contribution to monovinyl chlorophyll a (ng $L^{-1}$) from HPLC in CTD 114, 121, and 128. *Synechococcus* (SYN) values are from flow cytometry (see text). All other taxa are from Phytoclass assignments and represent prymnesiophytes (PRYM), pelagophytes (PELAG), diatoms type 2 (DIAT), dinoflagellates (A-DINO), prasinophytes group 3 (PRAS-3), prasinophytes group 1 (PRAS-1) and cryptophytes (CRYPT).

| Depth (m) | SYN | PRYM | PELAG | DIAT | A-DINO | PRAS-3 | PRAS-1 | CRYPT |
|---|---|---|---|---|---|---|---|---|
| **CTD 114** | | | | | | | | |
| 5 | 6.52 | 9.19 | 0.06 | 2.22 | 8.64 | 0.00 | 5.58 | 0.00 |
| 20 | 4.36 | 9.94 | 1.18 | 2.17 | 10.81 | 0.00 | 7.63 | 0.00 |
| 40 | 6.30 | 15.28 | 0.33 | 2.66 | 15.99 | 0.00 | 8.05 | 0.00 |
| 55 | 12.27 | 23.86 | 0.00 | 4.22 | 18.06 | 5.99 | 5.78 | 0.02 |
| 70 | 31.77 | 43.28 | 1.27 | 4.14 | 18.52 | 32.52 | 0.00 | 0.00 |
| 80 | 4.76 | 96.72 | 45.90 | 0.00 | 33.49 | 74.46 | 0.00 | 8.37 |
| **CTD 121** | | | | | | | | |
| 5 | 4.94 | 10.30 | 0.91 | 2.43 | 8.42 | 0.00 | 6.49 | 0.01 |
| 20 | 4.32 | 10.29 | 0.73 | 1.98 | 11.68 | 0.00 | 7.01 | 0.00 |
| 30 | 5.02 | 13.52 | 1.74 | 2.57 | 12.42 | 0.00 | 6.62 | 0.00 |
| 40 | 6.48 | 16.94 | 0.38 | 3.25 | 13.47 | 0.00 | 8.58 | 0.00 |
| 55 | 12.26 | 35.47 | 0.00 | 4.70 | 23.29 | 8.48 | 2.85 | 0.16 |
| 80 | 2.53 | 103.57 | 54.00 | 0.00 | 25.84 | 61.95 | 0.00 | 8.31 |
| **CTD 128** | | | | | | | | |
| 5 | 4.92 | 8.73 | 0.00 | 2.51 | 8.96 | 0.00 | 5.09 | 0.00 |
| 20 | 3.91 | 9.87 | 0.47 | 2.22 | 11.10 | 0.00 | 6.73 | 0.00 |
| 30 | 4.38 | 12.01 | 2.99 | 2.05 | 14.34 | 0.00 | 7.14 | 0.00 |
| 50 | 7.63 | 25.59 | 0.00 | 4.00 | 22.96 | 7.14 | 3.56 | 0.11 |
| 70 | 24.41 | 52.84 | 1.73 | 4.41 | 22.26 | 25.82 | 0.00 | 2.34 |
| 80 | 2.62 | 99.28 | 43.00 | 0.00 | 37.37 | 62.11 | 0.00 | 8.73 |

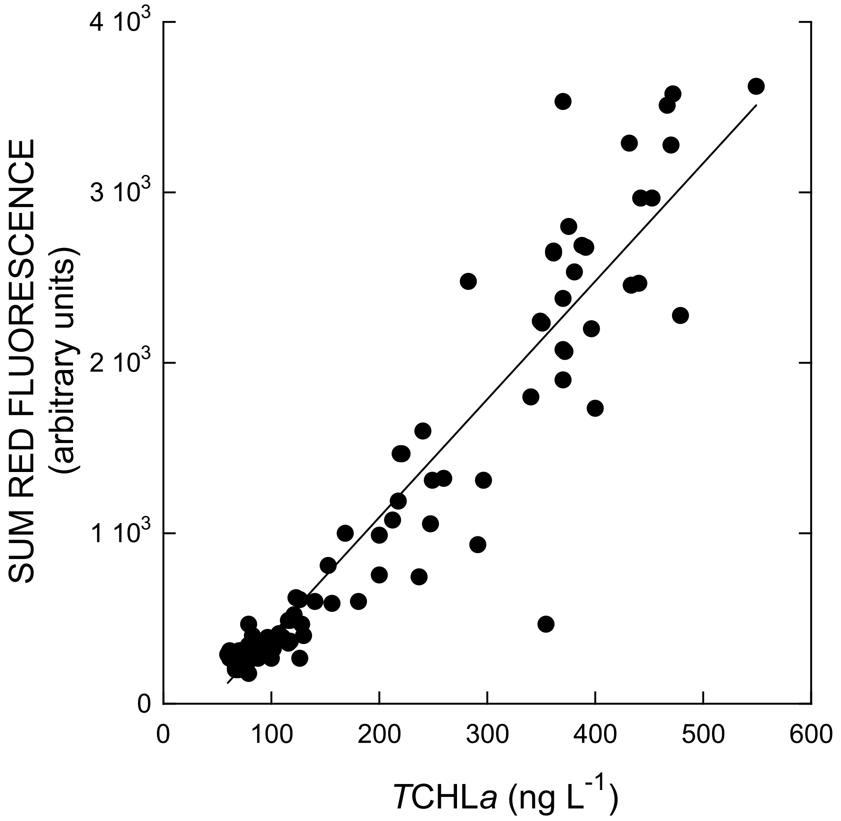


**Supplementary Figure S1.** Relationship between the sum of red fluorescence (SUM RF, abundance-weighted normalized chlorophyll fluorescence from flow cytometry) from all phytoplankton groups as a function of total CHLa (*T*CHL*a*, HPLC) in the same sample. Model II linear regression is: Sum RF = -289 + 6.93x*T*CHL*a*, $r^2 = 0.89$. For details on how Sum RF is estimated, see Selph et al., 2021.

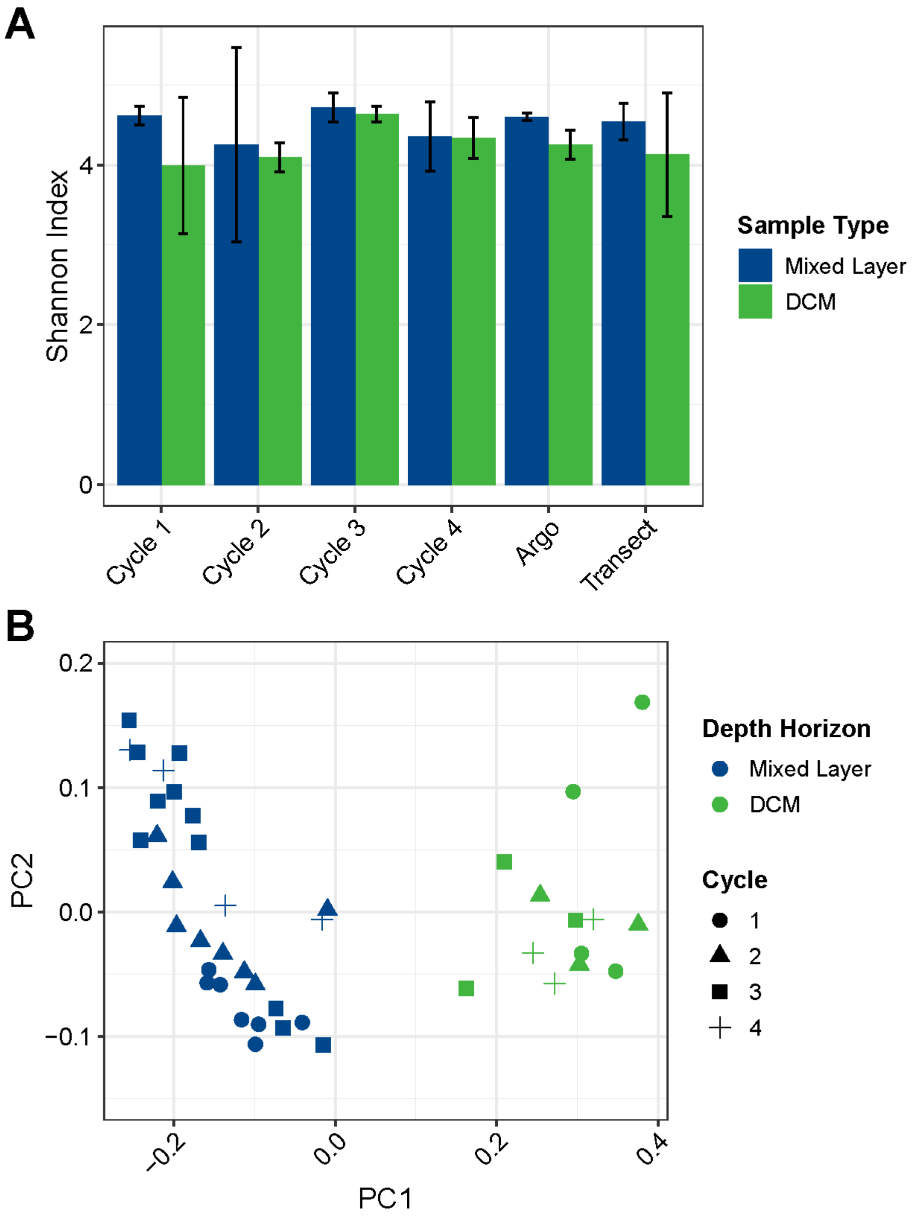


**Supplementary Figure S2**. Diversity of the eukaryotic phytoplankton ASVs (18S rRNA genes) between the mixed layer (blue) and DCM (green) among cycles, Argo stations, or transect stations. A) Alpha diversity expressed as the Shannon index. B) PCoA of Bray-Curtis dissimilarities.

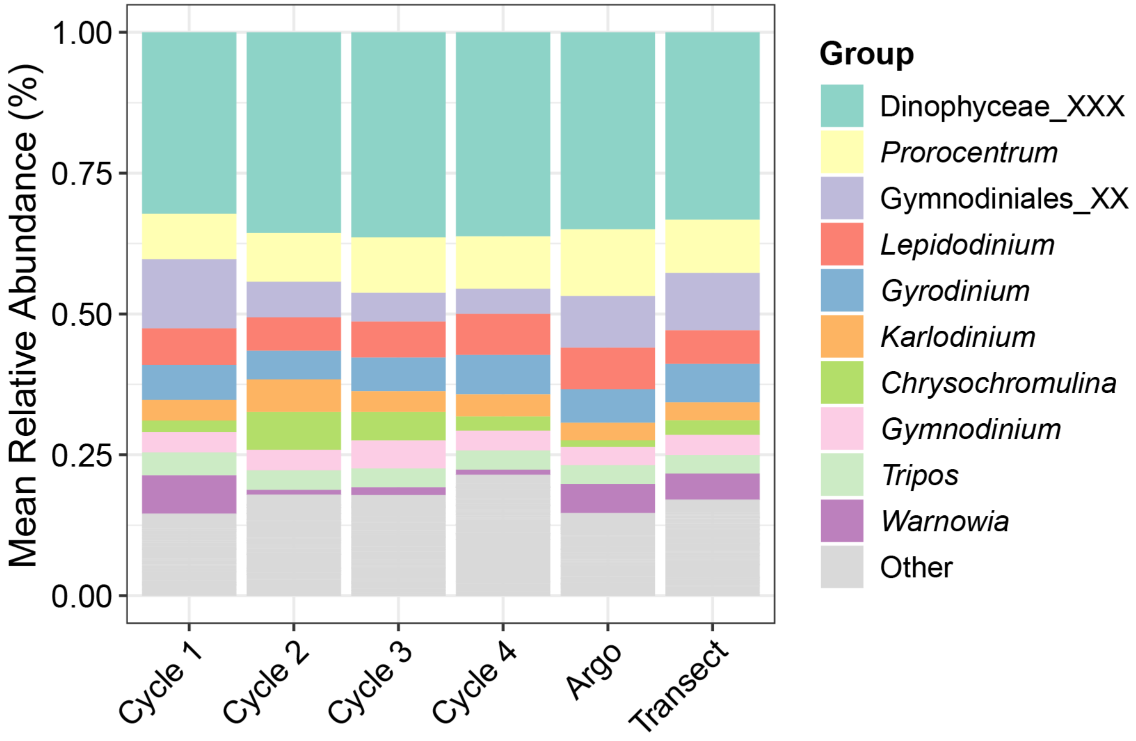


**Supplementary Figure S3.** Genus-level DNA-based relative abundances among A-DINO within each cycle, Argo stations, or transect samples. Where genus-level assignment is not available, the lowest taxonomic level is shown (*Gymnodiniales* or *Dinophyceae*).
